# An Engineered Contact Lens for Passive and Sustained Release of Lifitegrast, an Anti-Dry Eye Syndrome Drug

**DOI:** 10.1101/2021.04.10.439289

**Authors:** Changhua Mu, Victoria Lee, Yiran Liu, Ying Han, Gerard Marriott

**Affiliations:** Department of Bioengineering, University of California-Berkeley, Berkeley, California 94720, USA; Department of Ophthalmology, University of California, San Francisco, California 94158, USA

**Keywords:** Dry eye syndrome, Lifitegrast, Xiidra, contact lens, drug release

## Abstract

Lifitegrast is an FDA-approved drug that inhibits T-cell mediated inflammation associated with dry eye syndrome (DES). Lifitegrast is a potent inhibitor of the interaction between LFA-1 on T-cells and ICAM-1 on endothelial cells at the ocular surface. While effective in treating DES, 5% (81.2 mM) lifitegrast has low drug utilization and elicits off-target effects. Here we engineer contact lenses to release therapeutically-relevant doses of lifitegrast to every tear film for up to 10-hours. Lifitegrast is coupled to the polymer of the soft hydrogel lens via a photolabile (caged) crosslinker. Exposures of the lens to the 400-430 nm wavelengths of indoor daylight excite the caged crosslinker molecules and trigger a bond-cleavage reaction that releases authentic lifitegrast passively to the tear film. The photoproduct of the reaction remains chemically-linked to the polymer of the single-use lens. Our studies show that passive exposures of the lens to indoor light would generate an average of 990 nM lifitegrast to every tear film in a zero-order reaction for up to 10-hours. This concentration exceeds the *K*_d_ for the interaction between ICAM-1 and LFA-1 by ∼330-fold and would sustain inhibition of inflammatory responses at the ocular surface. The amount of lifitegrast released from the lens increases during exposures to outdoor sunlight. Over a 10-hour exposure to indoor light, a single lens would release 0.44% of the lifitegrast present in two drops of commercial 5% lifitegrast. Compared to tear-drop approaches, our engineered lenses would sustain the passive delivery of therapeutically-relevant doses of lifitegrast over a longer period, and exhibit improved drug utilization at a lower cost. Our technology could easily be integrated into daily-use contact lenses in order to prevent inflammation at the ocular surface, dry-eye and contact lens-mediated discomfort.

**Graphical Abstract:** 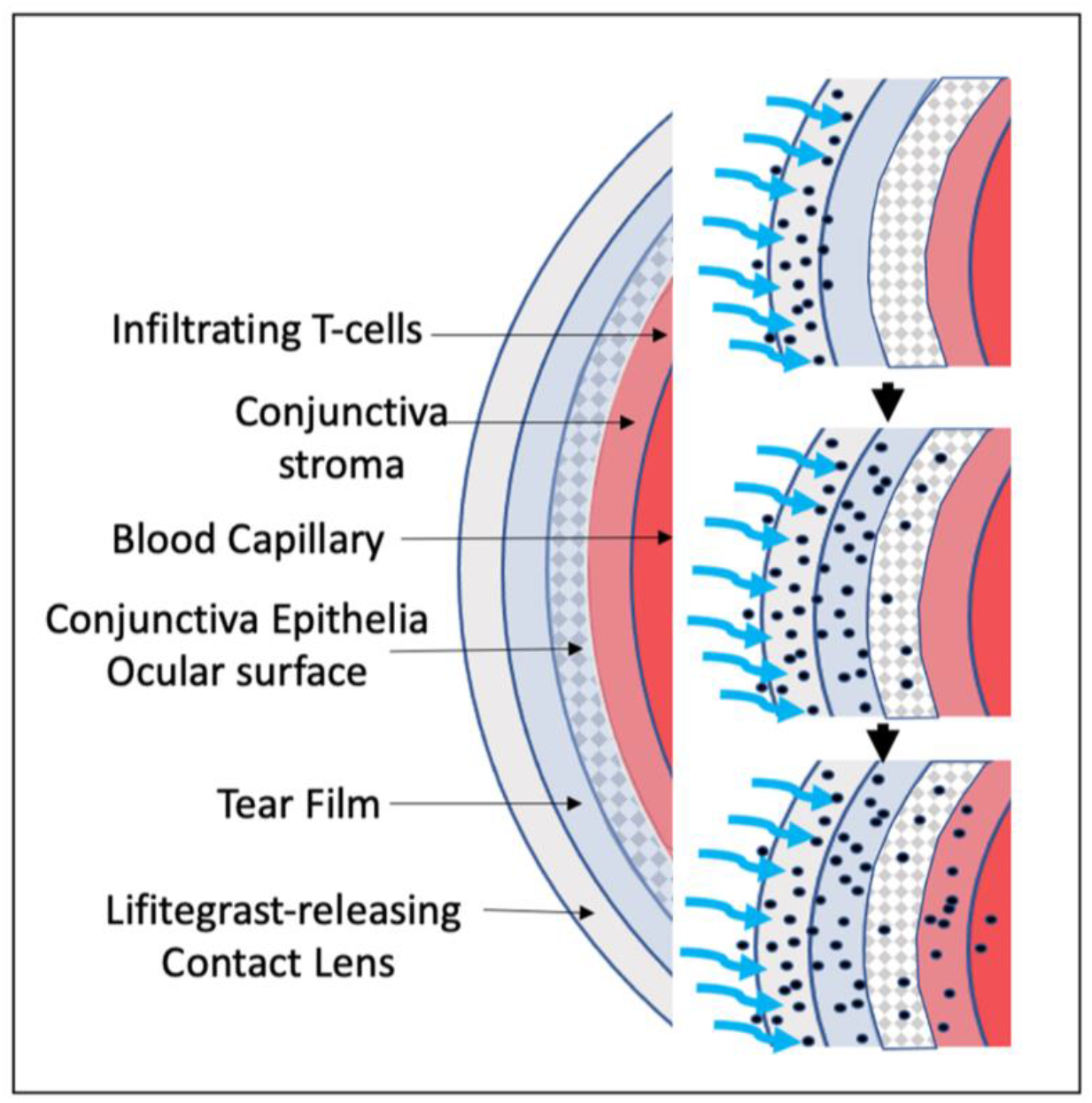

## INTRODUCTION

Approximately 40 million adults in the United States wear contact lenses^1^ with a significant number experiencing dry-eye syndrome (DES).^2^ DES symptoms in contact lens wearers include dryness, eye tiredness and soreness, irritation, scratchy sensations, blurry vision, and excessive blinking.^3^ These symptoms are typically managed by self-administration of artificial tears.^4^

DES is believed to result from T cell-mediated inflammation at the ocular surface and periocular tissue (Figure 1A,B).^5^ Inflammation at the ocular surface has been linked to the binding of lymphocyte function-associated antigen-1 (LFA-1) on T-cells, a heterodimeric protein of the integrin family,^6^ to the intercellular cell adhesion molecule-1 (ICAM-1) on conjunctiva epithelia. ICAM-1 is over-expressed on inflamed endothelial and epithelial cells, and on antigen-presenting cells (APCs).^7^ Formation of the LFA-1/ICAM-1 complex triggers the binding of the T-cell receptor (TCR) to the major histocompatibility complex (MHC) on the APC membrane to formi immunological synapses.^8^ Subsequent intracellular signaling events lead to the activation and proliferation of T-cells. Activated T-cells release inflammatory cytokines that may cause damage to ocular tissue, including the ocular surface.^9^ A competitive binding study has shown lifitegrast is a potent inhibitor of the interaction between LFA-1 and ICAM-1 (Figure 1C) on the surface of Jurkat T cells and HuT 78 T-cells, with an effective *K*_d_ of 3 nM.^10,11^

**Figure 1.**
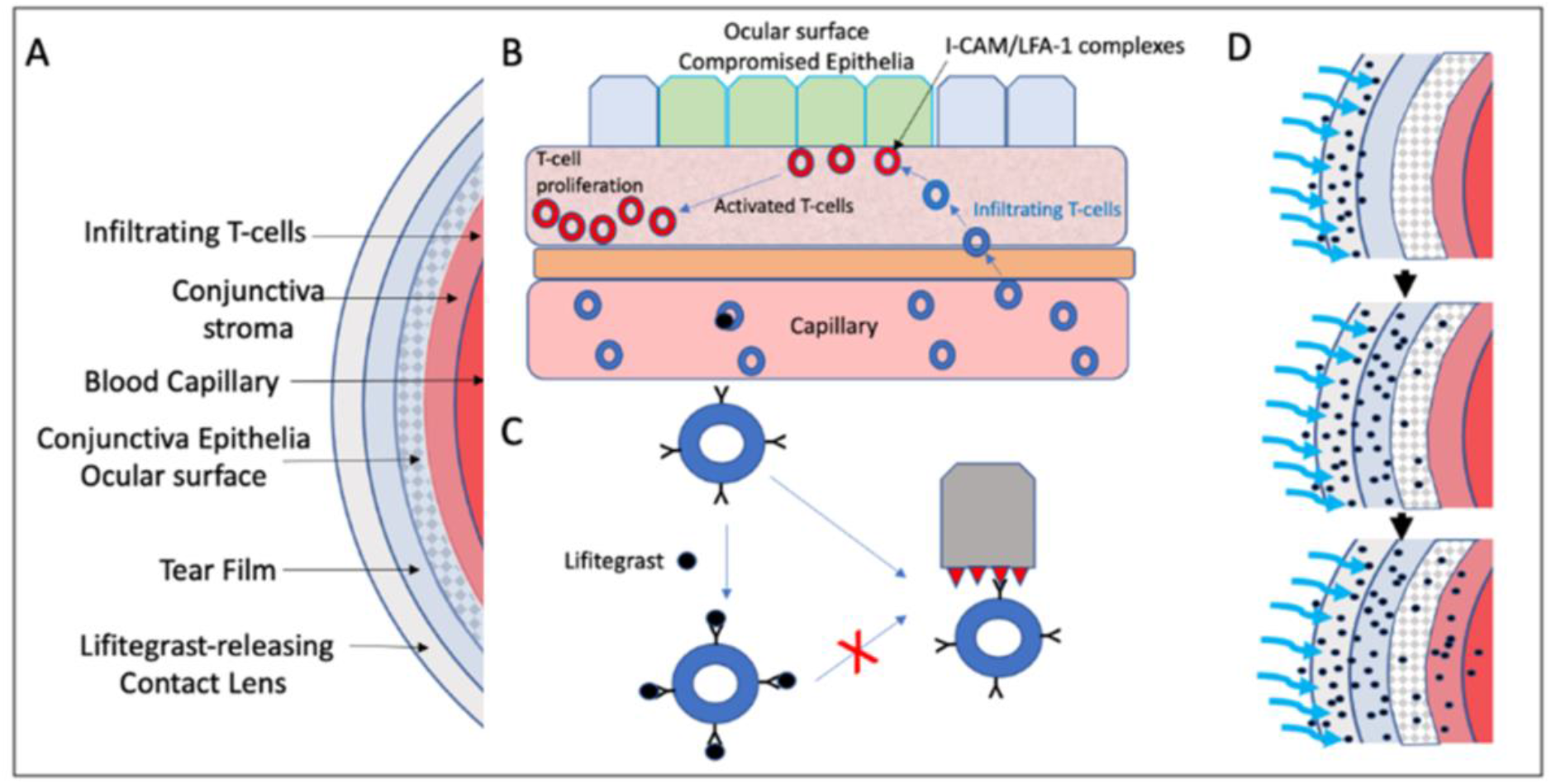
(A) Schematic representation of the engineered contact lens and the primary structure of the anterior eye. Damage to epithelia at the ocular surface results in the activation of T-cell mediated inflammatory responses; (B) The mechanism of action of lifitegrast: Stress or damage to epithelia at the ocular surface leads to the overexpression of ICAM-1; (C) Lifitegrast blocks the binding of LFA-1 to ICAM-1 on the surface of infiltrating T-cells. Inhibition of complex-formation reduces T-cell activation, formation of immunological synapses, release of cytokines, and further recruitment and proliferation of T-cells at the site of injury; (D) Our engineered contact lens release therapeutically-relevant doses of lifitegrast during exposures to indoor daylight (400-430 nm). The photo-released lifitegrast molecules diffuse progressively from the hydrogel of the lens to the tear film on the ocular surface, and to the conjunctiva stroma.

Lifitigrast has been shown to reduce T-cell activation and cytokine release and to mitigate downstream inflammatory processes (Figure. 1B).^12^ Lifitegrast is well-tolerated in patients, and provides rapid and sustained relief of DES compared to traditional treatments.^13-16^ Recently, the FDA approved a 5% lifitegrast ophthalmic solution (Xiidra™-Shire) as an eye-drop-based therapy for DES.^17,18^ Of note, the concentration of lifitegrast (in Xiidra) is 81.2 mM (50 mg/mL), which is 81,200 higher than the concentration required to treat severe cases of DES (1 µM).^12^ The high concentration of lifitegrast is necessary to overcome the low efficiency of drug delivery to inflamed tissue using eye-drops.^19^ The high concentration of lifitegrast also triggers off-target effects that include eye irritation, blurred vision, and dysgeusia (change in taste).^20^ Lifitegrast has also been used to manage DES in individuals with contact lens discomfort (CLD).^18^ In one study, lifitegrast was applied twice daily in eye-drop form using a protocol that required the individual to remove both contact lenses for 15 minutes, which is somewhat inconvenient.^18^ Previous studies have shown the difficulties among some patient groups in self-administering eye-drops, leading to additional expense and drug-wastage.^21^ Xiidra is already an expensive treatment for DES at ∼ $3.51/drop or $426.70 a month, and so efforts to reduce the cost of treatment would have a significant impact on healthcare costs.^22-24^

While the administration of 5% lifitegrast from teardrop applicators is effective in blocking the formation of LFA-1/ICAM-1 complexes at the ocular surface for a few hours, this drug-delivery method has low drug-utilization and elicits off-target effects.^12^ It has been reported that after dispensing an eye-drop, the concentration of lifitegrast in tear films is maintained at 1 µM for a few hours.^10,12,25^ Since the vast majority of drug molecules in the initial drop are rapidly cleared from eye, a smaller percentage must collect in local “reservoirs”, for example the conjunctiva stroma (Figure 1), from where lifitegrast is released to subsequent tear films.^10,12^ In this study, we introduce an engineered solution to sustain the release of lifitegrast to the tear film using contact lenses that release therapeutically-relevant doses (∼1 µM) to every tear film for up to 10 hours. Drug release is triggered from the lens during passive exposures to indoor and outdoor daylight (Figure 1D). We show that lens exposed to a constant intensity of indoor light release liftegrast to the bathing solution in a zero-order reaction. Herein we show our lenses would be more convenient in sustaining the release of therapeutically-relevant doses of lifitegrast than eye-drop approaches. They are also shown to have higher drug-utilization that would help to overcome overdosing and off-target effects.^20^ Finally, we highlight opportunities for the wearer to control the amount of drug released to the tear film, for example during exposures to outdoor sunlight.

## RESULTS AND DISCUSSION

An important aspect in the design of our contact lenses is to calculate how much caged lifitegrast must be coupled to the lens polymer to sustain release of therapeutic concentrations of lifitegrast to every tear film over the course of 10 hours. For this calculation, we assumed the individual blinks every 5 seconds^26^ and that each tear film has a volume of 5 mL.^27^ The dissociation constant (*K*_d_) for the lifitegrast/LFA-1 complex is related to the rate of complex dissociation (*k*_-1_), and the association rate of the complex (*k*_1_) by the expression:

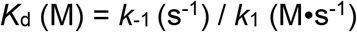

Using a *K*_d_ value of 3×10^−9^ M,^10,12^ and assuming a diffusion-controlled rate of binding of lifitegrast to LFA-1 in live cells of 10^7^ M^-1^s^-1^,^28^ we find lifitegrast molecules would dissociate from their LFA-1 complexes at an average rate (*k*_-1_) of 0.03 s^-1^. Thus, on exchanging the medium of cells whose LFA-1 membrane protein is stoichiometrically bound to lifitegrast, one would expect the drug to dissociate to 1/*e* of its initial concentration with a time constant of ∼33 seconds (1/k_-1_). In order to maintain the fully inhibited state, the medium (tear film) should be supplied with a steady-state concentration of lifitegrast at 30 nM (10 × *K*_d_). Next, we considered how much lifitegrast must be released from the lens to the tear film between successive blinks of the eye to maintain the 30 nM concentration. Assuming only 33% of lifitegrast molecules released from the inner and outer faces of the lens diffuses to the tear film during a 5-second inter-blink, then the lens itself should generate ∼100 nM lifitegrast in the between blinks or 5×10^−13^ mole or 3.08×10^−10^ g of lifitegrast (MW lifitegrast = 615.48 g·mol^−1^). Assuming uniform exposure of the central region of the contact lens to constant intensity daylight, the lens should release a total of 2217.6×10^−9^ g or ∼2.22 µg of lifitegrast over 7,200 blinks (10 hours). If we conservatively estimate that 10% of the caged lifitegrast is consumed from the lens over the 10-hour period, we would need to couple ∼22.2 µg of lifitegrast to each lens via the photolabile linker (or 36.3 µg of caged lifitegrast). Our optimized chemical-coupling protocol (shown later; Figure 2) is capable of linking 200 µg of caged lifitegrast to a single lens. Thus, we can ensure that contact lenses would have an adequate supply of caged lifitegrast to supply every tear film with 100 nM of the drug over a 10-hour period.

**Figure 2.**
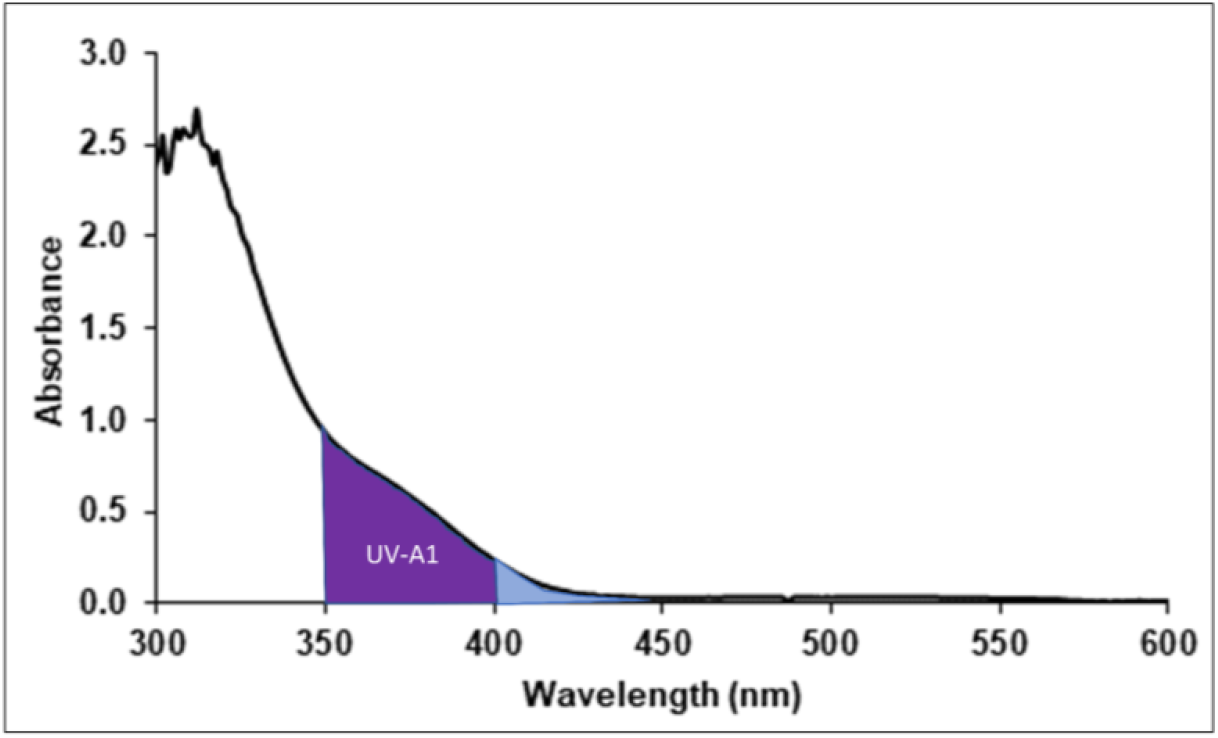
The absorption spectrum of the composite lens is dominated by the absorption of the caged crosslinker. Highlighted regions include the wavelength range of natural daylight for drug release (400∼430 nm), and the wavelength range associated with UV-A1 (340∼399 nm).

Lifitegrast is linked throughout the optically-transparent hydrogel polymer of the contact lens via a photolabile crosslinker. The caged crosslinker absorbs light from 300 nm to 430 nm, although we only consider photon interactions over the wavelength range of 400 – 430 nm (blue-violet). We note exposures of the eye to these blue-violet wavelengths is unavoidable, unless the individual wears sunglasses, or has UV-blue cut glass lenses.

### Coupling chemistry

We developed mild reaction conditions to couple lifitegrast to the caged crosslinker 4-[4-(1-hydroxyethyl)-2-methoxy-5-nitrophenoxy]butyric acid (hydroxyethyl photo-linker, or PL). All chemical reactions described herein were performed in a light-protected environment. First, we prepared the *N*-hydroxysuccinimide (NHS) ester at the carboxyl group of the crosslinker and then reacted with the amino group of 2-aminoethyl methacrylate to couple the methacrylate group (MA) to the photo-linker via an amide bond (**PL-MA**) (Reaction Scheme 1). Next, we brominated **PL-MA** with phosphorus tribromide (PBr_3_) to produce **Br-PL-MA**, which then reacted well with ammonia to generate the amine (**NH**_**2**_**-PL-MA**) (Reaction Scheme 2). In the last step, we used a standard carboxylic acid/amine coupling reaction using a mixture of catalysts, including *N, N*-diisopropylethylamine (DIPEA), 2-(1H-Benzotriazole-1-yl)-1,1,3,3-tetramethylaminium tetrafluoroborate (TBTU), and *N,N*-Dimethylpyridin-4-amine (DMAP), to form the amide bond with the carboxyl group of lifitegrast to cage lifitegrast (**LG-PL-MA**)(Reaction Scheme 2).^29^ Finally, LG-PL-MA is ready to co-polymerize with other monomers specified below through its MA group to fabricate the contact lens polymer (Reaction Scheme 3).

**Reaction Scheme 1.**
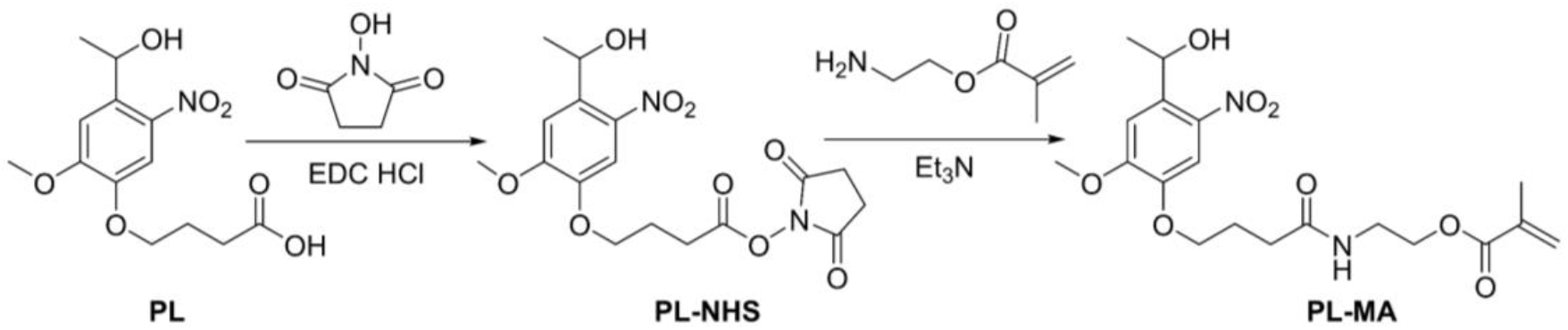
Preparation of methacrylate hydroxyethyl photolinker (PL-MA).

**Reaction Scheme 2.**
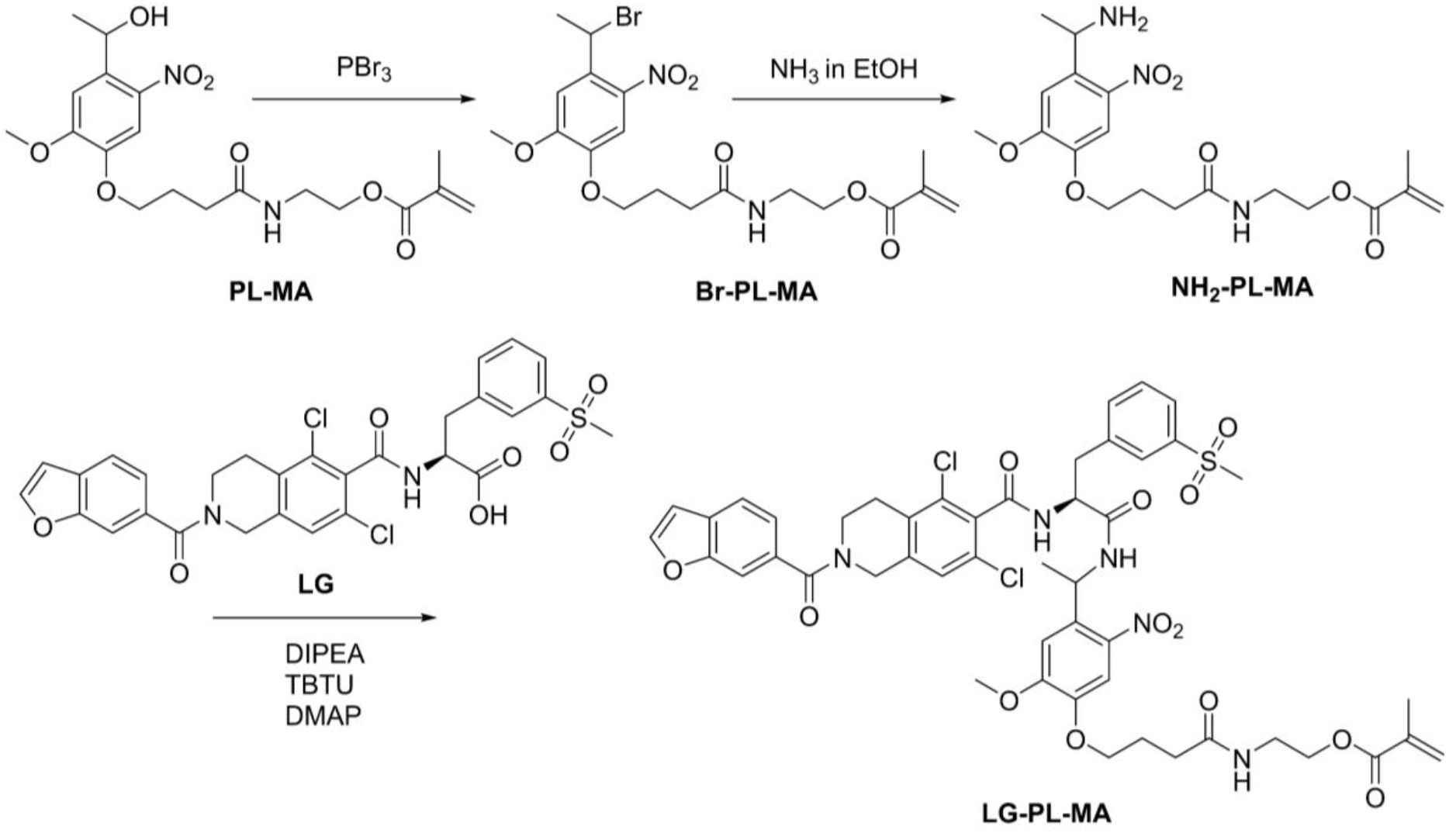
Preparation of caged lifitegrast (LG-PL-MA).

We used a CNC-milled template device to fabricate contact lenses. The volume sandwiched between the two components of the template was occupied with a pre-reaction mixture composed of caged lifitegrast (200 µg), poly-2-hydroxyethyl methacrylate (HEMA, 81 µL), *N,N*-dimethylallylamine (DMAA, 10 µL), methyl methacrylate (MMA, 1 µL), 4 Arm-PEG-Acrylate (8 µL of stock solution of 100 µg/µL in CH_2_Cl_2_) and 2,2-azobis(2-methylpropionitrile) (AIBN, 0.3 mg). Polymerization of this solution at 80 °C for 1 hour resulted in the formation of transparent hydrogels having the form and stiffness of a daily-use contact lens for humans.^30^ The functions of each component of the mixture are summarized as follows: HEMA was used as the backbone of the polymer; specific concentrations of DMAA and MMA were used to modulate the stiffness of the polymer; 4-Arm-PEG was used to increase the hydrophilicity of the polymer and helps to maintain water content. We employed AIBN as a heat-activated initiator. The dehydrated contact lens was washed and purified by exhaustive soakings in ethanol (8 × 4 mL) for 24 hours and hydrated in phosphate-buffered saline (PBS) (8 × 4 mL) for an additional 24-hour soaking. The lenses had a water content of 40% and were optically clear. Since the absorption of the photoprotection group crosslinker in the lens only extends to 430 nm, we would not expect it to impact color-perception, *i*.*e*., lenses are transparent over the visible range and indistinguishable in form and function to 2-hydroxyethyl methacrylate (HEMA)-based disposable contact lenses.^30^

**Reaction Scheme 3.**
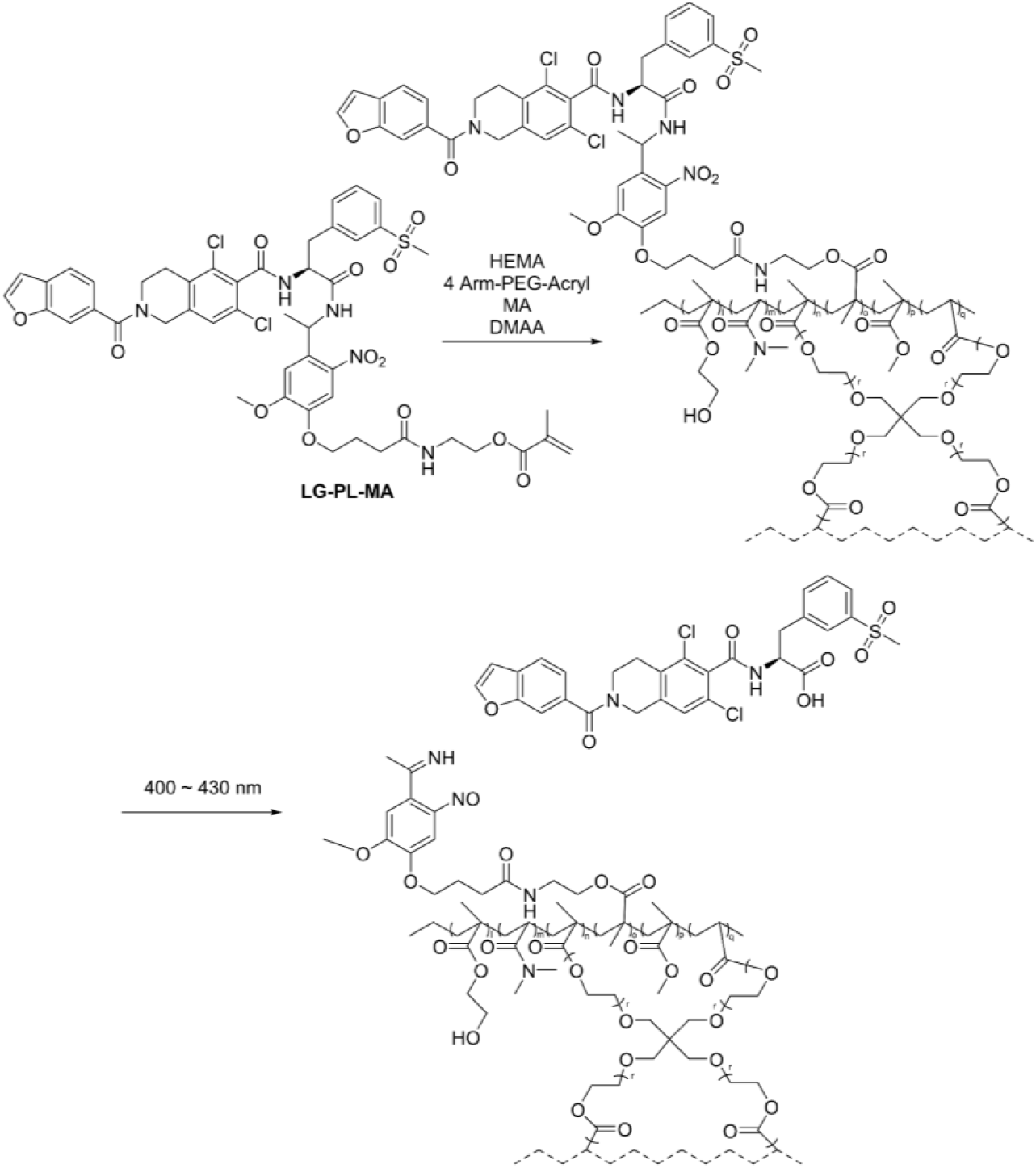
Co-polymerization of caged lifitegrast into contact lens hydrogel and subsequent release of authentic lifitegrast upon exposure to indoor daylight (400 – 430 nm).

### Spectroscopic characterization of our engineered contact lenses

The volume of a single contact lens is 56.6 µL (radius of 0.6 cm, thickness of 0.05 cm), and so the effective concentration of 200 µg (198.6 nmol) of caged lifitegrast chemically-coupled to the lens polymer is 3.58 mM. Using an average extinction coefficient (ε) of 1000 M^-1^cm^-1^ for the caged group between 400 – 430 nm,^30^ we uised the Beer-Lambert law to show caged groups in the lens would absorb ∼33.3% of all photons between 400 – 430 nm. Since the quantum yield for the uncaging reaction is 0.1 ^31,32^, then one equivalent of lifitegrast would be released from the lens after an average of 30 excitation events. Excitation of 4,5-dimethoxynitrobenzene (the caged group) between 400 – 430 nm results in rapid excited-state photoisomerization and cleavage of the amide bond that links liftegrast to the lens.^33^ We will show that exposure of the lens to 400∼430nm light generates free and authentic lifitegrast, carbon dioxide, and a 2-nitrosobenzaldehyde photoproduct, which remains attached to the lens polymer.^30^

Inspection of the absorption spectrum of the contact lens (Figure 2) measured using an Agilent diode array spectrophotometer,^19^ shows the caged group absorbs ∼×6 more photons over the UV-A1 range (340∼399 nm) compared to the 400∼430 nm range. The intensity of light over the 400∼430 nm range is greater for outdoor daylight compared to indoor daylight. Consequently, exposures to outdoor sunlight result in a higher rate of photon absorption by the caged crosslinker molecules in the lens that would increase the amount of lifitegrast released to the tear film. The same relationship would hold for the UV-A1 wavelengths, which are more intense on sunny days. Our contact lenses would respond to exposures of 340∼399nm photons in two important ways. First, the higher extinction coefficient of the caged group over 340∼399 nm (×3 - 6) would increase the amount of drug released from the lens. This property could be used to rapidly boost the concentration of lifitegrast in the tear film, for example during a bout of severe inflammation. Second, we have calculated caged crosslinker molecules in the lens would absorb ∼80% of the photons between 340∼399 nm,^34,35^ which would help protect the retina from UV-A1 photodamage. The lens would thereby provide three distinct and useful functions: First, to sustain the release of lifitegrast to each tear film for at least 10-hours; second, to provide corrective vision; and third, to act as a UV-A1 blocker.

### Characterization of light-mediated release of lifitegrast from contact lenses

Lifitegrast has a maximum UV-absorption band at 260 nm (Supporting Information, Figure S1), which we used to quantify the amount of drug released from the lens after exposures to blue-violet light (400 – 430 nm). We calculated the extinction coefficient of lifitegrast at 11300 ± 1400 M^-1^cm^-1^ from the intensity of absorption at 260 nm for known concentrations of the drug (n = 3) in PBS, (Supporting Information, Figure S2).

First, we investigated the kinetics of lifitegrast release from contact lenses during exposures to either 405 nm light delivered from an LED or indoor daylight. The protocols for these studies have been described by Mu *et al* (2018).^30^ The intensities of the 405 nm light and indoor daylight were measured using an Ohir meter manufactured by the Laser Measurement Group.^30^ Light delivered from a 395 nm LED (Cairn Research, UK) was directed through a Schott UG390 nm glass filter to remove all UV-A/B wavelengths. We measured the power of this transmitted light at 0.2 mW/cm^2^,^30^ and an absorption spectrophotometer to show a maximum wavelength of the filtered light was at ∼405 nm. Studies on the photo-release of lifitegrast from lenses using either 405 nm LED or natural indoor daylight were performed in triplicate (n = 3) as follows. First, a fully purified and hydrated lens (as described in the Methods section) was immersed in 2 mL of PBS in a scintillation vial that was completely transparent to each light source. We directed the 405 nm LED through a quartz lens (f = 10 cm) and located the vial at a distance to ensure complete and uniform illumination of the lens. We illuminated the lens in the vial for 1-minute, after which we gently agitated the vial for 15 minutes in the dark to flush the photo-released lifitegrast from the polymer. Next, we withdrew 1 mL of the bathing solution and recorded its UV-absorption spectrum using an Agilent 8453 diode array spectrophotometer (Figure 3A). After that, we returned the 1 ml volume to the mother solution in the vial and exposed the lens to the 405 nm LED for an additional minute. We repeated the same cycle of 405 nm exposure, stirring, and absorption measurement for 13-times. The absorption intensity of the withdrawn sample increased uniformly with illumination time, as shown in Figure 3A. The concentration of lifitegrast released from the lens at each time point was calculated from the absorption value at 260 nm using the measured extinction coefficient constant. The data plotted in Figure 3C shows the concentration of photo-released lifitegrast peaked at 25.0 µM after 13 minutes of accumulated exposures to 405 nm. This study shows that 50 nmol (30.77 µg) of lifitegrast was released from the lens to the bathing solution, which corresponds to an uncaging reaction rate (lifitegrast release rate) of 0.0758 nmol/s or 46.6 ng/s. The total amount of lifitegrast released from the lens represents 26.7% of the total caged lifitegrast coupled to the lens. Inspection of Figure 3C indicates the lens released free lifitegrast at a constant rate (zero-order release kinetics) in a 405 nm-dependent manner. We note devices that exhibit zeroth-order release kinetics are highly desired for long-term *in vivo* delivery of drugs.^36^

**Figure 3.**
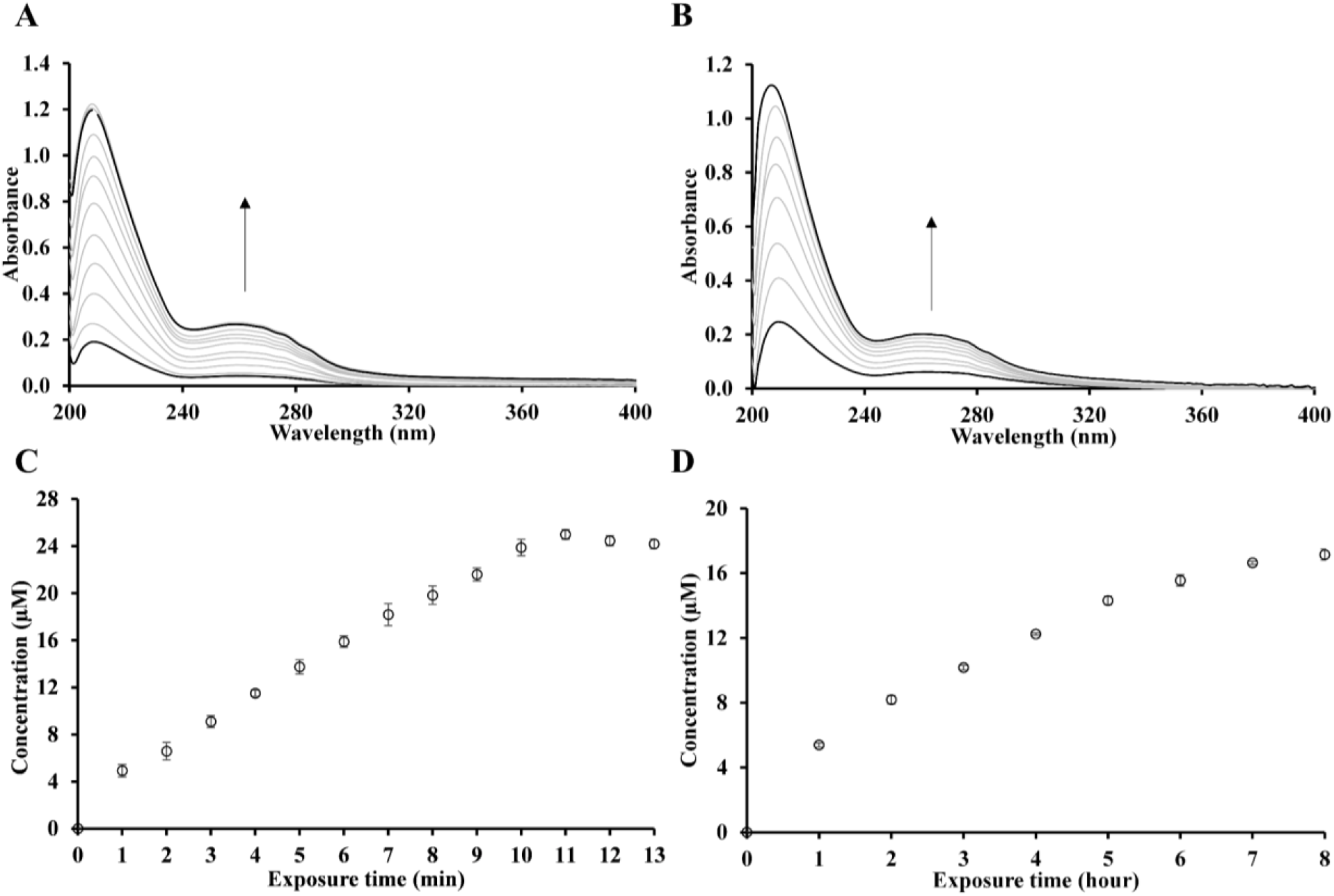
Representative UV-VIS spectra of the bathing solution of lenses exposed to (A) 405 nm LED light and (B) natural daylight, and their corresponding release kinetics (C) and (D) y-axis concentration data was calculated from the absorption value of lifitegrast at 260 nm.

Having shown that the lens generates free lifitegrast in the bathing solution to a concentration of 25.0 µM using the 405 nm LED source, we then conducted a second set of lifitegrast release studies using lenses exposed to natural indoor light. The contact lenses were processed as described above with the exception of being exposed to natural (indoor) daylight for 8 hours. These studies were performed from 8:00 am to 4:00 pm in December 2019 in a building on the campus of University of California, Berkeley. On these days, sunrise was at 7:15 am and sunset was at 4:50 pm.^37^ These studies were conducted in triplicate (n = 3). As shown in Figures 3B and 3D, we found that contact lenses released lifitegrast at an almost constant rate over the 8-hour exposure period. The average (cumulative) concentration of lifitegrast released from the lens to the bathing solution after 8 hours was measured at 17.16 μM, or 34.32 nmol (21.12 µg), which represents 68.6% of the maximum amount released at the 13-minute exposure to the more intense 405 nm light source.

By extrapolating the fitted line of data shown in Figure 3D to 10-hours of exposure to indoor daylight, we found our lenses would have released 35.66 nmol or 21.9 µg of lifitegrast, which corresponds to 19% of the caged lifitegrast we had crosslinked to the entire lens. The rate of lifitegrast release from the lens in this study was measured at 0.99 pmol/sec or 0.608 ng/sec, which corresponds to 990 nM in each tear film, or 330 × *K*_d_. It would seem likely that the concentration of lifitegrast released from the inner face of the lens to the interstitial fluid at the ocular surface would be higher than that in the volume of the tear film.

In summary, our results show that the amount of lifitegrast released to each tear film during passive exposures to indoor daylight over 10 hours would be sufficient to saturate the LFA-1 binding site on cells at the ocular surface, and thereby suppress T-cell mediated inflammatory responses associated with DES.

### Characterization of lifitegrast released from light-exposed contact lenses

We established lifitegrast photo-released from the lens to the bathing solution during exposures to 405 nm and indoor daylight was authentic by first recording and comparing the UV-vis absorption spectra of the bathing solution (Figure 3A and 3B) with a PBS solution of commercial lifitegrast (Supporting Information, Figure S1). The spectra were indistinguishable. We also conducted control studies (detailed below) to show that 260 nm absorbing species were not released from the lenses in the absence of light. Second, we evaporated the bathing solution of light-exposed lenses to dryness under vacuum, re-dissolved the residue in dimethyl sulfoxide-d6, and then recorded its 400 MHz ^1^H NMR spectrum (Figure 4). Analysis of the spectra shows the photo-released sample and authentic lifitegrast have identical resonance signals that coincide with reported values for lifitegrast.^38^ Importantly, we did not detect any resonances in the bathing solution that did not belong to lifitegrast, i.e., the bathing solution was free of impurities (Figure 4).

**Figure 4.**
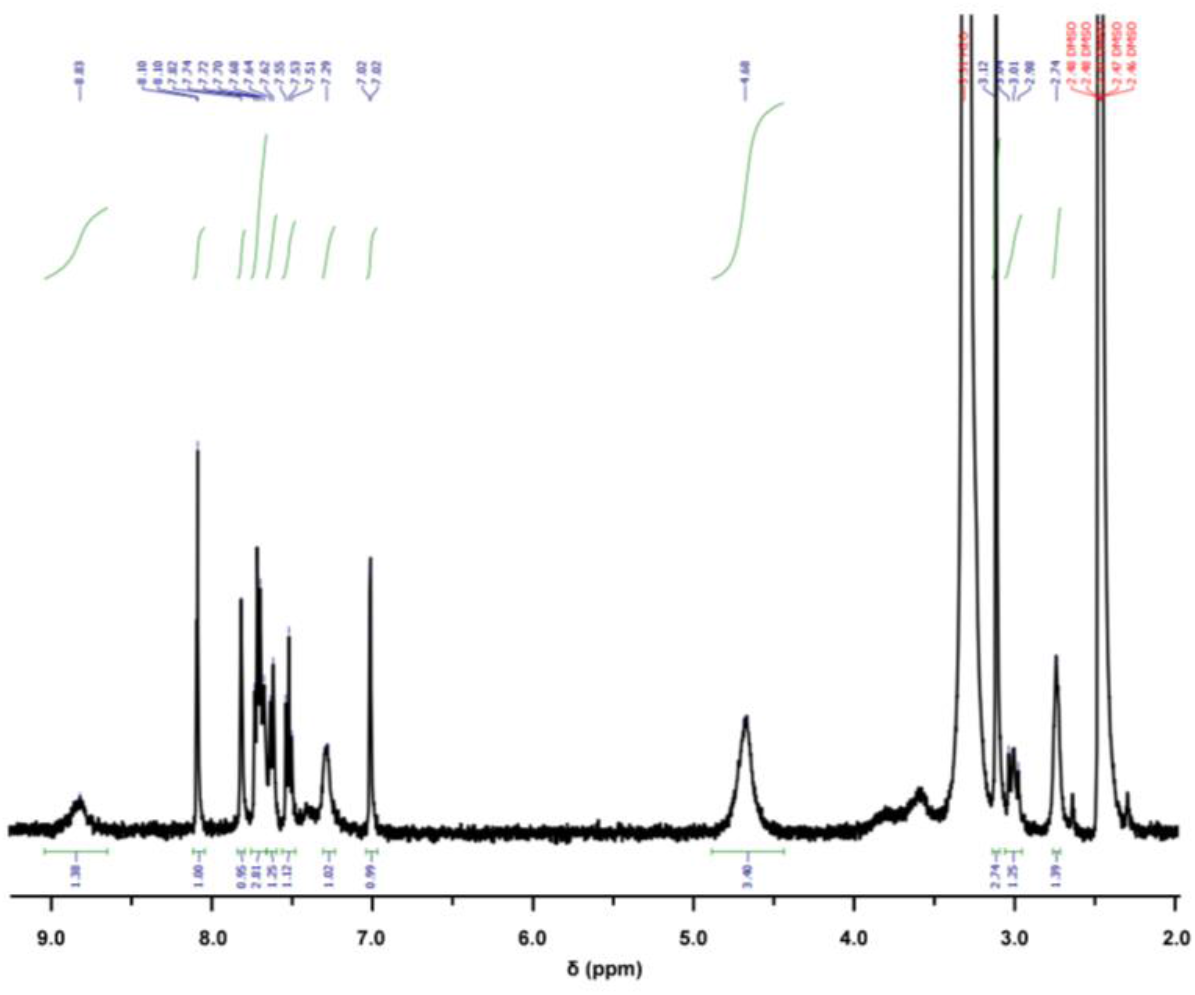
^1^H NMR (400 MHz, DMSO) spectrum of uncaged lifitegrast released from contact lenses after exposure to both 405 nm LED and indoor daylight. δ 8.83 (s, 1H), 8.10 (d, *J* = 2.2 Hz, 1H), 7.82 (s, 1H), 7.71 (q, *J* = 8.2 Hz, 3H), 7.63 (d, *J* = 7.8 Hz, 1H), 7.53 (t, *J* = 7.7 Hz, 1H), 7.29 (s, 1H), 7.02 (d, *J* = 2.1 Hz, 1H), 4.68 (s, 4H), 3.59 (s, 3H), 3.12 (s, 2H), 3.06 – 2.96 (m, 1H), 2.74 (s, 2H).

### Control studies

Control contact lenses without caged lifitegrast were prepared using the same fabrication method with the exception that **LG-PL-MA** was removed from the pre-reaction mixture. These control lenses were used to show our purification methods were effective in removing unreacted monomers after the polymerization reaction. This controlled study was necessary to prove the absorption signals recorded from the bathing solution at 260 nm arose exclusively from photo-released lifitegrast. In these experiments, dry lenses were fully hydrated by pre-soaking in 2 mL of fresh PBS on a shaker at a rate of 60 rpm for 4 hours in the dark. We showed that the lenses did not leach UV-absorbing components by recording absorption spectra of the bathing solution after 6 washes with PBS (Supporting Information, Figure S3). Next, we showed that the lenses did not leach any organic components by repeating this wash and measure procedure in ethanol. No measurable 260 nm absorption was recorded in either solvent after 6 washes.

In a second control study, we showed that uncaged lifitegrast was only released from the lens during exposures to the 405 nm LED, or natural indoor daylight. In these experiments, purified lenses coupled with caged lifitegrast were immersed in 2 mL of PBS on a shaker at a rate of 60 rpm in the dark for 3 days. Recordings of the absorption spectrum of the PBS solutions did not reveal any significant 260 nm absorption, which suggests that caged lifitegrast was not released in the absence of light (Supporting Information, Figure S4).

## CONCLUSIONS

In this study, we introduced a soft hydrogel contact lens that generates therapeutically-relevant concentrations of lifitegrast to tear films with zero-order release kinetics during passive exposures to indoor (and outdoor) daylight for up to 10 hours. These lenses offer significant advantages over the current eye-drop approaches to deliver lifitegrast that include passive release of therapeutically significant doses of the drug to every tear film for up to 10 hours at a constant (zero-order) rate of release, and the ability to control the amount of drug released from the lens during intermittent exposures to sunlight or to dim-light. A single lens releases 0.44% of the amount of lifitegrast present in 2-drops of Xiidra, which offers benefits in terms of cost, and reducing over-dosing and off-target effects. In addition to the drug release and vision correction functions, our lenses are highly effective in absorbing harmful UV-A1 wavelengths (340∼399 nm) and thereby act to protect the retina against exposures to harmful wavelengths. Finally, the lenses offer a simple, passive, and convenient means to manage DES-mediated contact lens discomfort among contact lens wearers.

## MATERIALS AND METHODS

### Materials

4-[4-(1-Hydroxyethyl)-2-methoxy-5-nitrophenoxy]butanoic acid (hydroxyethyl photo-linker, PL) was purchased from Novabiochem®. Lifitegrast was purchased from AChemBlock (Cat:L18012). 4 Arm-PEG-Acrylate (MW 10,000) was purchased from Laysan Bio, Inc.. 2-(1H-benzotriazole-1-yl)-1,1,3,3-tetramethylaminium tetrafluoroborate (TBTU), 4-(dimethylamino)pyridine (DMAP), *N,N*-diisopropylethylamine (DIPEA), 2-hydroxyethyl methacrylate (HEMA), methyl methacrylate (MMA), N,N-dimethylacrylamide (DMAA), 2,2’-azobis(2-methylpropionitrile) (AIBN), inhibitor remover for removing hydroquinone and monomethyl ether hydroquinone, and other chemical reagents were purchased from Sigma-Aldrich. No unexpected or unusually high safety hazards were encountered during our studies.

### Synthetic Procedures

All reactions were conducted in the dark, under N_2_ atmosphere protection, and with the rate of stirring set to 120 RPM. Unless specified, all experiments were conducted at room temperature.

#### Synthesis of PL-NHS

**PL-NHS** was synthesized using a previously reported synthesis.^30^

#### Synthesis of PL-MA

To a solution of **PL-NHS** (118.9 mg, 0.30 mmol) dissolved in 1.5 mL of DMF under stirring, a solution of 2-aminoethyl methacrylate hydrochloride (59.6 mg, 0.36 mmol) dissolved in 1.5 mL of methanol was added dropwise. After the solution was stirred for 15 minutes, triethylamine (125.4 μL, 91.1 mg, 0.90 mmol) was added. The reaction mixture was stirred at room temperature in the dark under a nitrogen gas atmosphere for 4 hours before it was diluted with 15 mL of chloroform and extracted with water (3 × 15 mL). The organic layer was dried over Na_2_SO_4_, filtered, and evaporated under vacuum to give a yellow oily product. The purity of the product was assured by TLC eluting with DCM/MeOH (9:1), showing only a single spot. Yield: 119.1 mg, 0.290 mmol, 96.7%. High-resolution electrospray ionization mass spectrometry (HR ESI-MS) of **PL-MA** m/z calculated for C_19_H_27_O_8_N_2_ ([M+H]^+^) 411.1762, found 411.1764, delta 0.50 ppm; m/z calculated for C_19_H_26_O_8_N_2_Na_1_ ([M+Na]^+^) 433.1581, found 433.1578, delta -0.78 ppm (Supporting Information, Figure S13).

#### Synthesis of Br-PL-MA

To a solution of **PL-MA** (119.1 mg, 0.290 mmol) in 6 mL of CH_2_Cl_2_, PBr_3_ (84.6 μL, 243.6 mg, 0.90 mmol) was added at 0 °C. After the reaction mixture was stirred overnight at room temperature in the dark under a nitrogen gas atmosphere, it was diluted with 15 mL of chloroform and extracted with water (3 × 15 mL). The organic layer was dried over Na_2_SO_4_, filtered, and evaporated under vacuum to produce a yellow solid. The product was used in the next step without further purification. HR ESI-MS of Br-PL-MA m/z calculated for C_19_H_25_O_7_N_2_Br_1_Na_1_ ([M+Na]^+^) 495.0737, found 495.0737, delta -0.07 ppm (Supporting Information, Figure S14).

#### Synthesis of NH2-PL-MA

To the above **Br-PL-MA** residual, 5 mL of ammonia solution (2 M in ethanol) was added. Excess ammonia and ethanol were removed under vacuum overnight while keeping the reaction mixture stirring at room temperature in the dark and under a nitrogen gas atmosphere. The residue was acidified with 5 mL of 0.5 M of HCl. The solution was then extracted with ethyl acetate (3 × 5 mL) to remove unreacted **Br-PL-MA** or other by-products. The aqueous phase solution was evaporated under vacuum to dryness, and 5 mL CH_2_Cl_2_ was added to the residue. The white insoluble suspension was filtered off and the filtrate was dried under vacuum to afford a yellow powder crude product. The crude product was purified by column chromatography with silica gel, eluting with a mixture of DCM/MeOH 9:1. The last band was collected, and the solvent was removed under vacuum to afford pure **NH**_**2**_**-PL-MA** as a pale yellow solid. Yield: 30.7 mg, 0.075 mmol, 25.9% from **PL-MA**. HR ESI-MS of **NH**_**2**_**-PL-MA** m/z calculated for C_19_H_28_O_7_N_3_ ([M+H]^+^) 410.1922, found 410.1918, delta -0.92 ppm (Supporting Information, Figure S15).

#### Synthesis of LG-PL-MA

**NH**_**2**_**-PL-MA** (30.7 mg, 0.075 mmol), lifitegrast (50.8 mg, 0.0825 mmol), DMAP (3.0 mg, 0.027 µmol), TBTU (28.9 mg, 0.09 mmol), and DIPEA (23.5 μL, 17.4 mg, and 0.135 mmol) were dissolved in 1.5 mL of DMF. After the resulting mixture was stirred for 16 hours at room temperature in the dark under a nitrogen gas atmosphere, 5 mL of 0.5 M of HCl was added and a pale-yellow precipitate formed immediately. The precipitate was collected, washed with 3 × 3 mL of water, and dissolved in 5 mL of CH_2_Cl_2_. The resulting solution was extracted with water (3 × 5 mL), dried over Na_2_SO_4_, and evaporated under vacuum to afford a yellow powder as the crude product. It was purified by column chromatography with silica gel eluting with a mixture of CH_2_Cl_2_/MeOH (9.5: 0.5, v/v). The second from the last band (R_f_ = 0.2) was collected and evaporated under vacuum to afford pure **LG-PL-MA** as a pale yellow solid. Yield: 20.8 mg, 0.0207 mmol, 27.6%. HR ESI-MS of **LG-PL-MA** m/z calculated for C_48_H_50_O_13_N_5_Cl_2_S_1_ ([M+H]^+^) 1006.2497, found 1006.2501, delta 0.36 ppm; m/z calculated for C_48_H_49_O_13_N_5_Cl_2_Na_1_S_1_ ([M+Na]^+^) 1028.2317, found 1028.2318, delta 0.11 ppm (Supporting Information, Figure S16).

### Preparation of Lifitegrast Release Contact Lenses

The transparent mock contact lens was fabricated using a method adapted from a literature report with materials used for making functional contact lenses for human use._39_ To a mixture of monomers composed of HEMA (810 µL), MMA (10 µL), and DMAA (100 µL), ∼10 inhibitor remover beads were added, shaken well for 30 minutes, and filtered off. A solution of 4 Arm-PEG-Acrylate (8 mg dissolved in 80 µL CH_2_Cl_2_), photo-initiator AIBN (3 mg), and **LG-PL-MA** (2 mg) was added to the mixture of the monomers and mixed well. The stock solution was ready to be used for fabricating the lifitegrast release contact lenses.

The stock solution (100 µL) was added to the 2-component case mold, which had been coated with Gel Slick solution (Lonza, cat. No.:50640). The mold was heated at 80°C for 1 hour to get a dry contact lens, which was purified by soaking in ethanol (6 × 3 mL) for 24 hours and then in PBS (6 × 3 mL) for another 24 hours.

### Measurement of Extinction Coefficient of Lifitegrast

Pure lifitegrast (Advanced ChemBlocks, Cat: L18012) was measured with a calibrated high accuracy and precision analytical balance (Model: Mettler Toledo xs205) and dissolved in 1.0 mL of PBS solution measured by a calibrated pipette. Lifitegrast has two absorptions at 205 and 260 nm. The absorption at 260 nm was used to determine the extinction coefficient since 205 nm is too close to the instrument’s shortest detectable wavelengths (∼200 nm), potentially introducing noise and errors in the data.

Absorption, measured at 260 nm, was recorded for a series of lifitegrast solutions prepared via a serial dilution from the stock solution (4:5 dilutions). Using the Beer-Lambert Law, experimental absorption values, and known concentrations of lifitegrast, the molar extinction coefficient constant was calculated. The experiments were performed in triplicate (n = 3) and the average value and standard deviations were calculated (Supporting Information, Figure S2).

### UV-vis Test Contact Lenses Coupled with Caged Lifitegrast

To inspect the UV-vis absorption of the lens coupled with caged lifitegrast, we used the above-mentioned method but using flat molds made of silicone isolator between two the glass slides to prepare the flat lenses (diameter 13 mm, thickness 0.5 mm) coupled with caged lifitegrast. We also made blank lenses without coupling caged lifitegrast and used them as the baseline control. We measured the UV-vis absorption spectrum as shown in Figure 2.

### Spectroscopic Studies of the Light-Mediated Release of Lifitegrast from Contact Lens

A contact lens loaded with caged lifitegrast was immersed in 2 mL of PBS in a 20 mL scintillation glass vial equipped with a small stir bar. Two types of light sources were used to trigger the release of lifitegrast from the contact lens to its bathing solution: 405 nm light from a 395 nm LED source (Cairn Research, UK) filtered through a Schott UG390 nm filter to remove the UV-A (<400 nm; power = 0.2 mW/cm^2^ measured using an Ohir meter manufactured by Laser Measurement Group), and natural indoor daylight on typical rainy winter days (December 2019) (n = 3) from 8 am to 4 pm (sunrise at 7:15 am and sunset at 4:50 pm) on the University of California, Berkeley campus. The release process of photo-uncaged lifitegrast from the contact lens to its bathing solution was monitored using UV-vis absorption spectroscopy recorded by an Agilent 8453 UV–vis spectrophotometer (Agilent Technologies).

In the 405 nm LED light-mediated release experiments, the lenses were directly exposed to 405 nm LED light for 1 minute from a distance of approximately 30 cm and stirred at a rate of 60 rpm in the dark for 15 minutes to allow the full dissociation of uncaged lifitegrast into the bathing solution. A UV-vis absorption spectrum of the bathing solution was recorded each time until no further change of the spectra was observed. In the indoor daylight-mediated release experiments, the contact lenses were exposed to normal indoor daylight inside a glass vial while stirred at a rate of 60 rpm. The absorption spectra of the bathing solution were recorded every hour over an exposure period of 8 hours. In both experiments, the vials were capped when possible to prevent the evaporation of water from the PBS bathing solution. Absorptions of the bathing solutions at 260 nm were used for the release kinetic studies.

## ASSOCIATED CONTENT

### Supporting Information

The Supporting Information includes follows:

Additional data and figures including instrument and experiment photographs, high-resolution electrospray ionization mass spectroscopic spectra, UV−vis absorption study.

## Author Information

### Author Contributions

C.M. designed the experimental protocols, synthesized, and characterized the desired compounds, fabricated the functional contact lens, collected and analyzed data, and wrote the manuscript. V.L. assisted in chemical synthesis, compound purification, data collection, and made the molds of the 2-component case to fabricate the contact lenses. Y.L. assisted in the hydrogel system studies, contact lens preparation, and data collection. Y.H. guided and helped in the initial development of the contact lens, and participated in the writing and editing of the manuscript. G.M. proposed the study, directed the experiments, analyzed critical data, wrote and edited the manuscript with input from all other authors. All authors read, discussed, and corrected the manuscript.

### Notes

The authors declare the following competing financial interest(s): Work detailed in the manuscript has been submitted to the USPTO as a provisional patent.

## ACKNOWLEDGMENTS

The authors acknowledge the research funds supported by the Tsinghua-Berkeley Shenzhen Institute. The authors acknowledge QB3/Chemistry Mass Spectrometry Facility at the University of California Berkeley for running mass spec and providing data. The authors also acknowledge Jacobs Makerspace for access to equipment and materials to fabricate the 2-component case mold for preparing mock contact lenses.

## REFERENCES

1 Cope, J. R. et al.. Risk Behaviors for Contact Lens-Related Eye Infections Among Adults and Adolescents - United States, 2016. MMWR Morb Mortal Wkly Rep 66, 841–845 (2017).

2 Nichols, J. J. et al.. The TFOS International Workshop on Contact Lens Discomfort: executive summary. Invest Ophthalmol Vis Sci 54, TFOS7–TFOS13 (2013).

3 The definition and classification of dry eye disease: report of the Definition and Classification Subcommittee of the International Dry Eye WorkShop (2007). Ocul Surf 5, 75–92 (2007).

4 Papas, E. B. et al.. The TFOS International Workshop on Contact Lens Discomfort: report of the management and therapy subcommittee. Invest Ophthalmol Vis Sci 54, TFOS183–203 (2013).

5 Stevenson, W., Chauhan, S. K. & Dana, R. Dry eye disease: an immune-mediated ocular surface disorder. Arch. Ophthalmol. 130, 90–100, doi:10.1001/archophthalmol.2011.364 (2012).

6 Kuerzinger, K. & Springer, T. A. Purification and structural characterization of LFA-1, a lymphocyte function-associated antigen, and Mac-1, a related macrophage differentiation antigen associated with the type three complement receptor. J. Biol. Chem. 257, 12412–12418 (1982).

7 Marlin, S. D. & Springer, T. A. Purified intercellular adhesion molecule-1 (ICAM-1) is a ligand for lymphocyte function-associated antigen 1 (LFA-1). Cell (Cambridge, Mass.) 51, 813–819, doi:10.1016/0092-8674(87)90104-8 (1987).

8 Grakoui, A. et al.. The immunological synapse: A molecular machine controlling T cell activation. Science. 285, 221–227, doi:10.1126/science.285.5425.221 (1999).

9 Hogg, N., Laschinger, M., Giles, K. & McDowall, A. T-cell integrins: More than just sticking points. J. Cell Sci. 116, 4695–4705, doi:10.1242/jcs.00876 (2003).

10 Murphy, C. J. et al.. The pharmacologic assessment of a novel lymphocyte function-associated antigen-1 antagonist (SAR 1118) for the treatment of keratoconjunctivitis sicca in dogs. Invest. Ophthalmol. Visual Sci. 52, 3174–3180, doi:10.1167/iovs.09-5078 (2011).

11 Zhong, M. et al.. Discovery and Development of Potent LFA-1/ICAM-1 Antagonist SAR 1118 as an Ophthalmic Solution for Treating Dry Eye. ACS Med. Chem. Lett. 3, 203–206, doi:10.1021/ml2002482 (2012).

12 Semba, C. P. & Gadek, T. R. Development of lifitegrast: a novel T-cell inhibitor for the treatment of dry eye disease. Clin. Ophthalmol. 10, 1083–1094, doi:10.2147/OPTH.S110557 (2016).

13 Zhong, M. et al.. Discovery of tetrahydroisoquinoline (THIQ) derivatives as potent and orally bioavailable LFA-1/ICAM-1 antagonists. Bioorg. Med. Chem. Lett. 20, 5269–5273, doi:10.1016/j.bmcl.2010.06.145 (2010).

14 Zhong, M. et al.. Structure-activity relationship (SAR) of the α-amino acid residue of potent tetrahydroisoquinoline (THIQ)-derived LFA-1/ICAM-1 antagonists. Bioorg. Med. Chem. Lett. 21, 307–310, doi:10.1016/j.bmcl.2010.11.014 (2011).

15 1 Gadek, T. R. et al.. Generation of an LFA-1 antagonist by the transfer of the ICAM-1 immunoregulatory epitope to a small molecule. Science (Washington, DC, U. S.) 295, 1086–1089, doi:10.1126/science.295.5557.1086 (2002).

16 Keating, S. M. et al.. Competition between intercellular adhesion molecule-1 and a small-molecule antagonist for a common binding site on the α1 subunit of lymphocyte function-associated antigen-1. Protein Sci. 15, 290–303, doi:10.1110/ps.051583406 (2006).

17 Keating, G. M. Lifitegrast Ophthalmic Solution 5%: A Review in Dry Eye Disease. Drugs 77, 201–208 (2017).

18 Gonzalez, A. L. Safety and efficacy of lifitegrast 5% ophthalmic solution in contact lens discomfort. Clin Ophthalmol 12, 2079–2085 (2018).

19 Haber, S. L. et al.. Lifitegrast: a novel drug for patients with dry eye disease. Ther Adv Ophthalmol 11, 2515841419870366 (2019).

20 Donnenfeld, E. D. et al.. Safety of Lifitegrast Ophthalmic Solution 5.0% in Patients With Dry Eye Disease: A 1-Year, Multicenter, Randomized, Placebo-Controlled Study. Cornea 35, 741–748 (2016).

21 Morrison, P. W. J. & Khutoryanskiy, V. V. Advances in ophthalmic drug delivery. Ther. Delivery 5, 1297–1315, doi:10.4155/tde.14.75 (2014).

22 Lifitegrast (Xiidra) for Dry Eye Disease. JAMA 317, 1473–1474 (2017).

23 Donnenfeld, E. D., Perry, H. D., Nattis, A. S. & Rosenberg, E. D. Lifitegrast for the treatment of dry eye disease in adults. Expert Opin. Pharmacother. 18, 1517–1524, doi:10.1080/14656566.2017.1372748 (2017).

24 Allen, M. Drug Companies Make Eyedrops Too Big, And You Pay For The Waste, Shots Health News From NPR. https://www.npr.org/sections/health-shots/2017/10/18/558358137/drug-companies-make-eyedrops-too-big-and-you-pay-for-the-waste October 18, 2017).

25 Dustin, M. L. & Groves, J. T. Receptor signaling clusters in the immune synapse. Annu. Rev. Biophys. 41, 543–556, doi:10.1146/annurev-biophys-042910-155238 (2012).

26 Craig, J. P., Tomlinson, A. & McCann, L. in Encyclopedia of the Eye (ed Darlene A. Dartt) 254–262 (Academic Press, 2010).

27 Scherz, W., Doane, M. G. & Dohlman, C. H. Tear volume in normal eyes and keratoconjunctivitis sicca. Albrecht Von Graefes Arch Klin Exp Ophthalmol 192, 141–150 (1974).

28 Erickson, J., Goldstein, B., Holowka, D. & Baird, B. The effect of receptor density on the forward rate constant for binding of ligands to cell surface receptors. Biophys. J. 52, 657–662, doi:10.1016/S0006-3495(87)83258-7 (1987).

29 El-Faham, A. & Albericio, F. Peptide Coupling Reagents, More than a Letter Soup. Chem. Rev. 111, 6557–6602, doi:10.1021/cr100048w (2011).

30 Mu, C., Shi, M., Liu, P., Chen, L. & Marriott, G. Daylight-Mediated, Passive, and Sustained Release of the Glaucoma Drug Timolol from a Contact Lens. ACS Cent. Sci. 4, 1677–1687, doi:10.1021/acscentsci.8b00641 (2018).

31 Ottl, J., Gabriel, D. & Marriott, G. Preparation and Photoactivation of Caged Fluorophores and Caged Proteins Using a New Class of Heterobifunctional, Photocleavable Crosslinking Reagents. Bioconjugate Chem. 9, 143–151, doi:10.1021/BC970147O (1998).

32 Marriott, G., Roy, P. & Jacobson, K. Preparation and light-directed activation of caged proteins. Methods Enzymol. 360, 274–288 (2003).

33 Marriott, G. Caged Protein Conjugates and Light-Directed Generation of Protein Activity: Preparation, Photoactivation, and Spectroscopic Characterization of Caged G-Actin Conjugates. Biochemistry 33, 9092–9097, doi:10.1021/bi00197a010 (1994).

34 Ottl, J., Gabriel, D. & Marriott, G. Preparation and Photoactivation of Caged Fluorophores and Caged Proteins Using a New Class of Heterobifunctional, Photocleavable Crosslinking Reagents. Bioconjugate Chem. 9, 143–151, doi:10.1021/bc970147o (1998).

35 Marriott, G., Roy, P. & Jacobson, K. Preparation and light-directed activation of caged proteins. Methods Enzymol. 360, 274–288 (2003).

36 Ali, M. et al.. Zero-order therapeutic release from imprinted hydrogel contact lenses within in vitro physiological ocular tear flow. J. Controlled Release 124, 154–162, doi:10.1016/j.jconrel.2007.09.006 (2007).

37 Berkeley, California, USA — Sunrise, Sunset, and Daylength, December 2019, <https://www.timeanddate.com/sun/usa/berkeley?month=12&year=2019> (2019).

38 Organic Spectroscopy International Lifitegrast, SAR 1118, <https://orgspectroscopyint.blogspot.com/2014/09/lifitegrast-sar-1118.html> (2014).

39 Lai, C.-F., Li, J.-S., Fang, Y.-T., Chien, C.-J. & Lee, C.-H. UV and blue-light antireflective structurally colored contact lenses based on a copolymer hydrogel with amorphous array nanostructures. RSC Adv. 8, 4006–4013, doi:10.1039/c7ra12753g (2018).

