## supplementary information for "An Engineered Contact Lens for Passive and Sustained Release of Lifitegrast, an Anti-Dry Eye Syndrome Drug"

**Table of Contents**

(1) UV-vis measurement of molar extinction coefficient of lifitegrast…..………….……….S2

(2) UV-vis measurement of the control experiments…..…………………………………....S3

(3) UV-vis measurement of 405 nm LED light mediated lifitegrast release…....…….…....S5

(4) UV-vis absorption spectra of daylight mediated lifitegrast release…..…….………......S7

(5) High-resolution electrospray ionization mass spectrometry…..…….…………….….S12

(6) UV-vis absorption spectra of synthetic compounds…..…………………..……….…..S16

### **(1) UV-vis Measurement of Molar Extinction Coefficient of Lifitegrast**

| Wavelength (nm) | Absorbance |
| --- | --- |
| 190 | 0.245007 |
| 191 | 0.292167 |
| 192 | 0.279461 |
| 193 | 0.321439 |
| 194 | 0.378057 |
| 195 | 0.398344 |
| 196 | 0.399516 |
| 197 | 0.4217 |
| 198 | 0.490563 |
| 199 | 0.508524 |
| 200 | 0.533246 |
| 201 | 0.650141 |
| 202 | 0.778056 |
| 203 | 0.81532 |
| 204 | 0.820879 |
| 205 | 0.817114 |
| 206 | 0.812356 |
| 207 | 0.807044 |
| 208 | 0.799942 |
| 209 | 0.790292 |
| 210 | 0.77665 |
| 211 | 0.759777 |
| 212 | 0.739177 |
| 213 | 0.715636 |
| 214 | 0.69001 |
| 215 | 0.663352 |
| 216 | 0.636209 |
| 217 | 0.609691 |
| 218 | 0.583161 |
| 219 | 0.557395 |
| 220 | 0.531527 |
| 221 | 0.506295 |
| 222 | 0.48123 |
| 223 | 0.456648 |
| 224 | 0.432677 |
| 225 | 0.408638 |
| 226 | 0.38464 |
| 227 | 0.36024 |
| 228 | 0.336175 |
| 229 | 0.313333 |
| 230 | 0.290641 |
| 231 | 0.268337 |
| 232 | 0.245209 |
| 233 | 0.221908 |
| 234 | 0.199753 |
| 235 | 0.1794 |
| 236 | 0.162487 |
| 237 | 0.149041 |
| 238 | 0.138895 |
| 239 | 0.13193 |
| 240 | 0.127142 |
| 241 | 0.124197 |
| 242 | 0.122777 |
| 243 | 0.122583 |
| 244 | 0.123336 |
| 245 | 0.124715 |
| 246 | 0.126636 |
| 247 | 0.129249 |
| 248 | 0.132043 |
| 249 | 0.134961 |
| 250 | 0.137786 |
| 251 | 0.140709 |
| 252 | 0.143802 |
| 253 | 0.146556 |
| 254 | 0.149292 |
| 255 | 0.151807 |
| 256 | 0.153714 |
| 257 | 0.154988 |
| 258 | 0.155704 |
| 259 | 0.156016 |
| 260 | 0.156167 |
| 261 | 0.155898 |
| 262 | 0.155065 |
| 263 | 0.153666 |
| 264 | 0.151726 |
| 265 | 0.149708 |
| 266 | 0.147823 |
| 267 | 0.146132 |
| 268 | 0.144298 |
| 269 | 0.141886 |
| 270 | 0.138323 |
| 271 | 0.133481 |
| 272 | 0.128327 |
| 273 | 0.124039 |
| 274 | 0.120807 |
| 275 | 0.118251 |
| 276 | 0.11573 |
| 277 | 0.111938 |
| 278 | 0.105943 |
| 279 | 0.098351 |
| 280 | 0.090158 |
| 281 | 0.0826135 |
| 282 | 0.0768909 |
| 283 | 0.0728841 |
| 284 | 0.0691786 |
| 285 | 0.0645833 |
| 286 | 0.0585189 |
| 287 | 0.0520558 |
| 288 | 0.0456057 |
| 289 | 0.0394416 |
| 290 | 0.0336761 |
| 291 | 0.028379 |
| 292 | 0.023663 |
| 293 | 0.0196362 |
| 294 | 0.016192 |
| 295 | 0.0133252 |
| 296 | 0.0110245 |
| 297 | 0.00922441 |
| 298 | 0.00771952 |
| 299 | 0.00650024 |
| 300 | 0.00549078 |
| 301 | 0.00469875 |
| 302 | 0.00427723 |
| 303 | 0.00378275 |
| 304 | 0.00334263 |
| 305 | 0.00317287 |
| 306 | 0.00286436 |
| 307 | 0.00248384 |
| 308 | 0.00241327 |
| 309 | 0.00221872 |
| 310 | 0.002285 |
| 311 | 0.00214911 |
| 312 | 0.00199223 |
| 313 | 0.00195312 |
| 314 | 0.00191736 |
| 315 | 0.00185585 |
| 316 | 0.00190067 |
| 317 | 0.00177193 |
| 318 | 0.00177574 |
| 319 | 0.00183105 |
| 320 | 0.00189924 |
| 321 | 0.00187254 |
| 322 | 0.00170946 |
| 323 | 0.00175905 |
| 324 | 0.00182104 |
| 325 | 0.00182438 |
| 326 | 0.00181103 |
| 327 | 0.00184155 |
| 328 | 0.00179291 |
| 329 | 0.00172901 |
| 330 | 0.00173044 |
| 331 | 0.00176096 |
| 332 | 0.00167799 |
| 333 | 0.0017457 |
| 334 | 0.00175142 |
| 335 | 0.0016942 |
| 336 | 0.00171995 |
| 337 | 0.00171614 |
| 338 | 0.00171661 |
| 339 | 0.00170422 |
| 340 | 0.00168371 |
| 341 | 0.00170994 |
| 342 | 0.00178528 |
| 343 | 0.00180292 |
| 344 | 0.00185442 |
| 345 | 0.00179672 |
| 346 | 0.00179625 |
| 347 | 0.00182009 |
| 348 | 0.00176096 |
| 349 | 0.00180721 |
| 350 | 0.00176668 |
| 351 | 0.00182867 |
| 352 | 0.00177336 |
| 353 | 0.0017252 |
| 354 | 0.00175619 |
| 355 | 0.0018754 |
| 356 | 0.00191545 |
| 357 | 0.00165844 |
| 358 | 0.00192881 |
| 359 | 0.00214243 |
| 360 | 0.00204945 |
| 361 | 0.00196505 |
| 362 | 0.00154924 |
| 363 | 0.00133085 |
| 364 | 0.00159645 |
| 365 | 0.00169802 |
| 366 | 0.0016222 |
| 367 | 0.00169897 |
| 368 | 0.00192595 |
| 369 | 0.00190401 |
| 370 | 0.00178576 |
| 371 | 0.00172377 |
| 372 | 0.00177479 |
| 373 | 0.00183487 |
| 374 | 0.00175953 |
| 375 | 0.0015769 |
| 376 | 0.00173235 |
| 377 | 0.0018425 |
| 378 | 0.00183344 |
| 379 | 0.00180197 |
| 380 | 0.00184202 |
| 381 | 0.00163889 |
| 382 | 0.00194597 |
| 383 | 0.00175238 |
| 384 | 0.00185299 |
| 385 | 0.0017519 |
| 386 | 0.00170803 |
| 387 | 0.00189686 |
| 388 | 0.00195408 |
| 389 | 0.00182247 |

Figure S1. UV-vis absorption spectrum of lifitegrast in PBS with two absorption peaks at 210 and 260 nm.

| Concentration (M) | Absorbance |
| --- | --- |
| 0.00010648 | 1.11 |
| 8.51839e-05 | 0.89115 |
| 6.81471e-05 | 0.72924 |
| 5.45177e-05 | 0.61901 |
| 4.36141e-05 | 0.50321 |
| 3.48913e-05 | 0.42017 |
| 2.79131e-05 | 0.36009 |
| 2.23304e-05 | 0.28039 |
| 1.78644e-05 | 0.22291 |
| 1.42915e-05 | 0.17897 |
| 1.14332e-05 | 0.14152 |
| 9.14655e-06 | 0.11809 |

| Concentration (M) | Absorbance |
| --- | --- |
| 0.00010648 | 1.2321 |
| 8.51839e-05 | 1.0065 |
| 6.81471e-05 | 0.77574 |
| 5.45177e-05 | 0.66353 |
| 4.36141e-05 | 0.54024 |
| 3.48913e-05 | 0.45961 |
| 2.79131e-05 | 0.35838 |
| 2.23304e-05 | 0.30028 |
| 1.78644e-05 | 0.25918 |
| 1.42915e-05 | 0.20987 |
| 1.14332e-05 | 0.18924 |
| 9.14655e-06 | 0.15045 |

| Concentration (M) | Absorbance |
| --- | --- |
| 0.00010648 | 1.3814 |
| 8.51839e-05 | 1.1837 |
| 6.81471e-05 | 0.99598 |
| 5.45177e-05 | 0.78297 |
| 4.36141e-05 | 0.63268 |
| 3.48913e-05 | 0.51319 |
| 2.79131e-05 | 0.43282 |
| 2.23304e-05 | 0.3694 |
| 1.78644e-05 | 0.28693 |
| 1.42915e-05 | 0.23228 |
| 1.14332e-05 | 0.18849 |
| 9.14655e-06 | 0.16786 |

Figure S2. Plots of UV-vis absorptions of lifitegrast at 260 nm vs. its standard solutions. The experiment was repeated three times (n = 3). The extinction coefficient of lifitegrast at 260 nm was determined as 11300 ± 1400 M^-1^ cm^-1^.

### **(2)** **UV-vis Measurement of the Control Experiments**

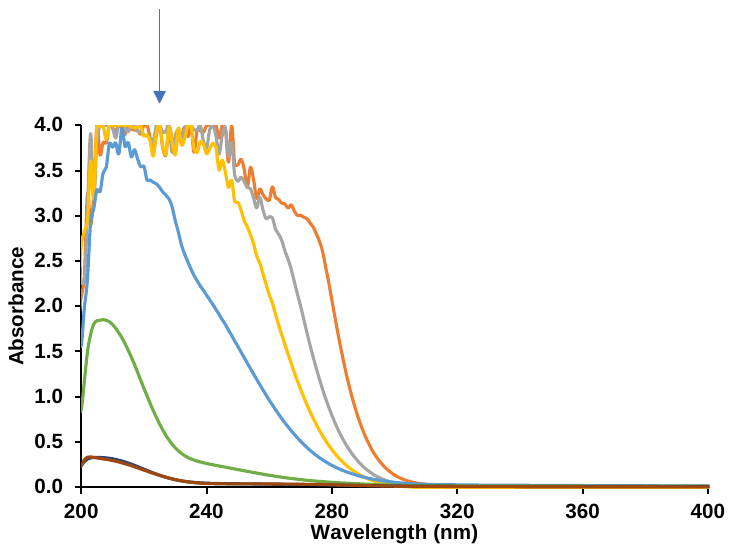

Figure S3. UV-vis absorption spectra showing the process of purifying a contact lens by washing it with PBS over time in the dark. In each measurement, the lens was soaked in 2 mL of fresh PBS solution for 4 hours, and the UV-vis absorption of the bathing solution was recorded. The procedure was repeated until no signal was observed, suggesting 6 changes of 2 mL of PBS over the course of 6 hours is enough to purify the contact lens.

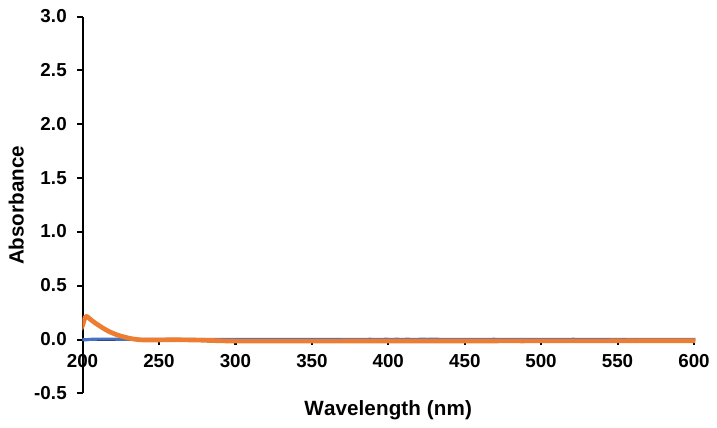

Figure S4. UV-vis absorption spectrum of the 2-mL bathing solution, in which a lens coupled with caged lifitegrast was immersed for 3 days in the dark. No absorption intensity corresponding to caged lifitegrast, lifitegrast, and the photoproduct was recorded.

### **(3) UV-vis Measurement of 405 nm LED Light Mediated Lifitegrast Release**

**a**

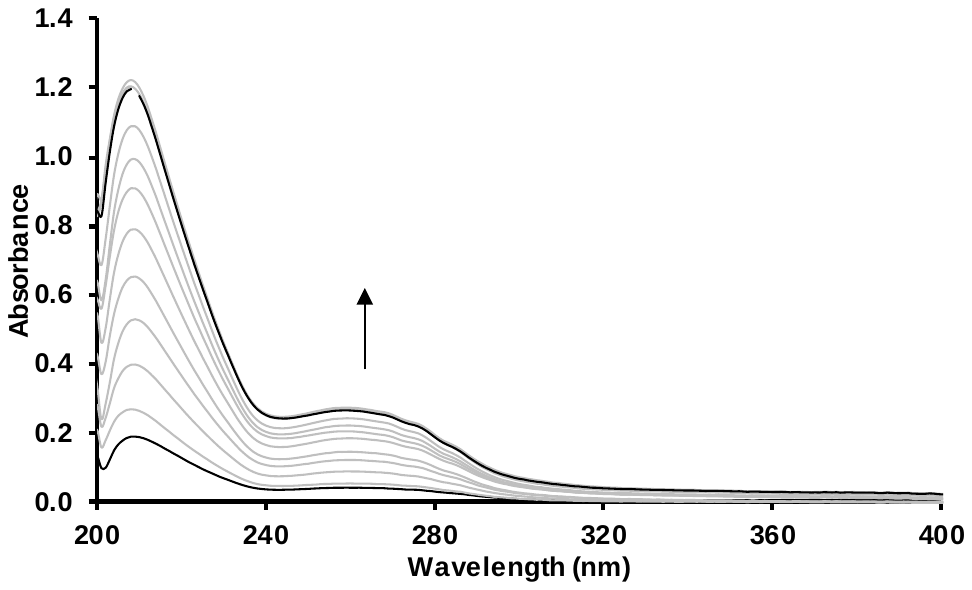

**b**

| Exposure time (mins) | Absorbance |
| --- | --- |
| 0 | 0 |
| 1 | 0.0425892 |
| 2 | 0.0550613 |
| 3 | 0.0897732 |
| 4 | 0.122885 |
| 5 | 0.146896 |
| 6 | 0.185947 |
| 7 | 0.20582 |
| 8 | 0.222708 |
| 9 | 0.243565 |
| 10 | 0.270317 |
| 11 | 0.274164 |
| 12 | 0.268244 |
| 13 | 0.266115 |

Figure S5. 405 nm LED light-mediated lifitegrast release from a contact lens (Trial - 1). (a) A lifitegrast release contact lens was immersed in 2 mL of PBS. At each measurement, the sample was exposed to 405 nm LED light for 1 minute and stirred in the dark at a rate of 60 rpm for 15 minutes. 1 mL of the bathing solution was withdrawn, and then returned to the mother solution after its UV-vis spectra was recorded, (b) plot of the absorption at 260 nm versus time.

**a**

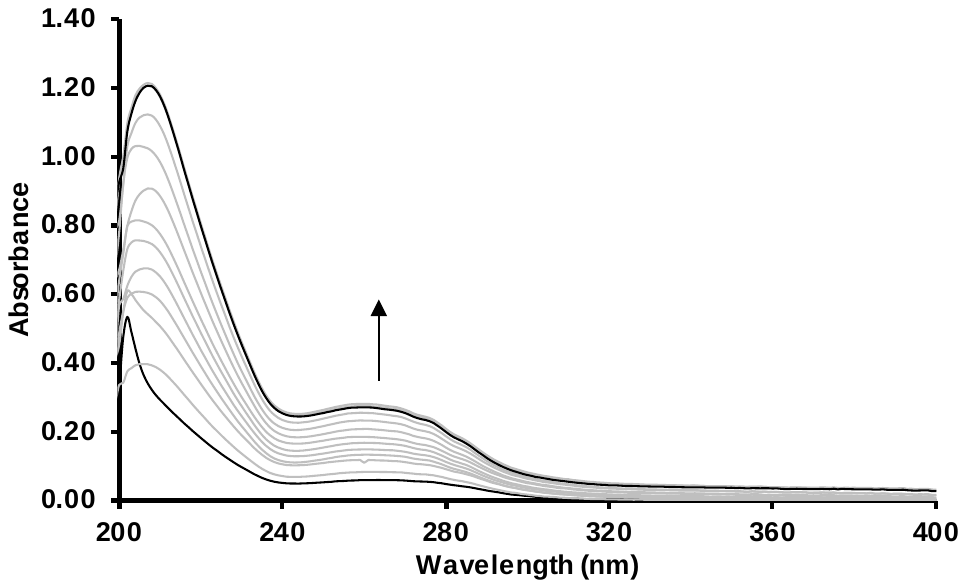

**b**

| Exposure time (mins) | Absorbance |
| --- | --- |
| 0 | 0 |
| 1 | 0.0582027 |
| 2 | 0.0821743 |
| 3 | 0.10893 |
| 4 | 0.132315 |
| 5 | 0.148016 |
| 6 | 0.166969 |
| 7 | 0.184122 |
| 8 | 0.207176 |
| 9 | 0.23159 |
| 10 | 0.254118 |
| 11 | 0.280146 |
| 12 | 0.274208 |
| 13 | 0.270037 |

Figure S6. 405 nm LED light-mediated lifitegrast release from a contact lens (Trial - 2). (a) a lifitegrast release contact lens was immersed in 2 mL of PBS. At each measurement, the sample was exposed to 405 nm LED light for 1 minute and stirred in the dark at a rate of 60 rpm for 15 minutes. 1 mL of the bathing solution was withdrawn, and then returned to the mother solution after its UV-vis spectra was recorded, (b) plot of the absorption at 260 nm versus time.

### **(4) UV-vis Measurement of Indoor Daylight Mediated Lifitegrast Release**

**a**

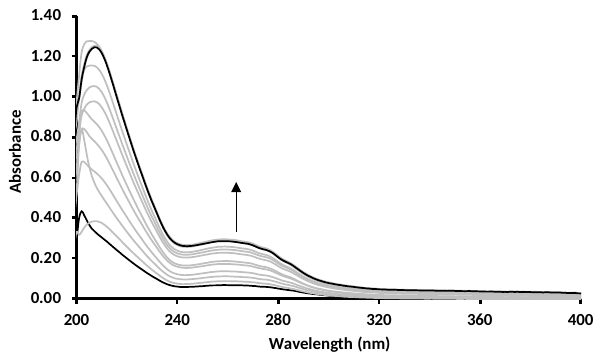

**b**

| Exposure time (mins) | Absorbance |
| --- | --- |
| 0 | 0 |
| 1 | 0.0659828 |
| 2 | 0.0860901 |
| 3 | 0.109621 |
| 4 | 0.134872 |
| 5 | 0.171011 |
| 6 | 0.185398 |
| 7 | 0.226423 |
| 8 | 0.242283 |
| 9 | 0.256704 |
| 10 | 0.285145 |
| 11 | 0.292789 |
| 12 | 0.2869 |
| 13 | 0.283716 |

Figure S7. 405 nm LED light-mediated lifitegrast release from a contact lens (Trial - 3). (a) a lifitegrast release contact lens was immersed in 2 mL of PBS. At each measurement, the sample was exposed to 405 nm LED light for 1 minute and stirred in the dark at a rate of 60 rpm for 15 minutes. 1 mL of the bathing solution was withdrawn, and then returned to the mother solution after its UV-vis spectra was recorded, (b) plot of the absorption at 260 nm versus time.

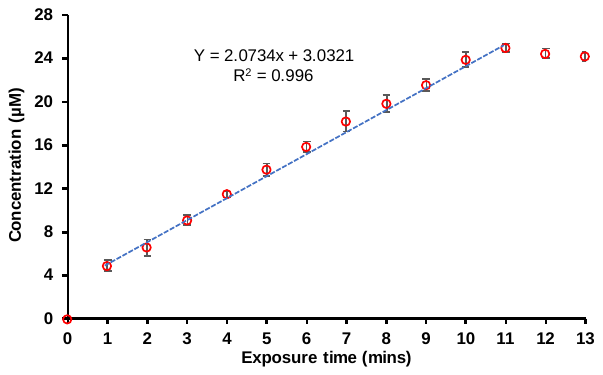

Figure S8. A graph of 405 nm LED light-mediated lifitegrast release (n = 3) from a contact lenses, including the average and standard deviation.

**a**

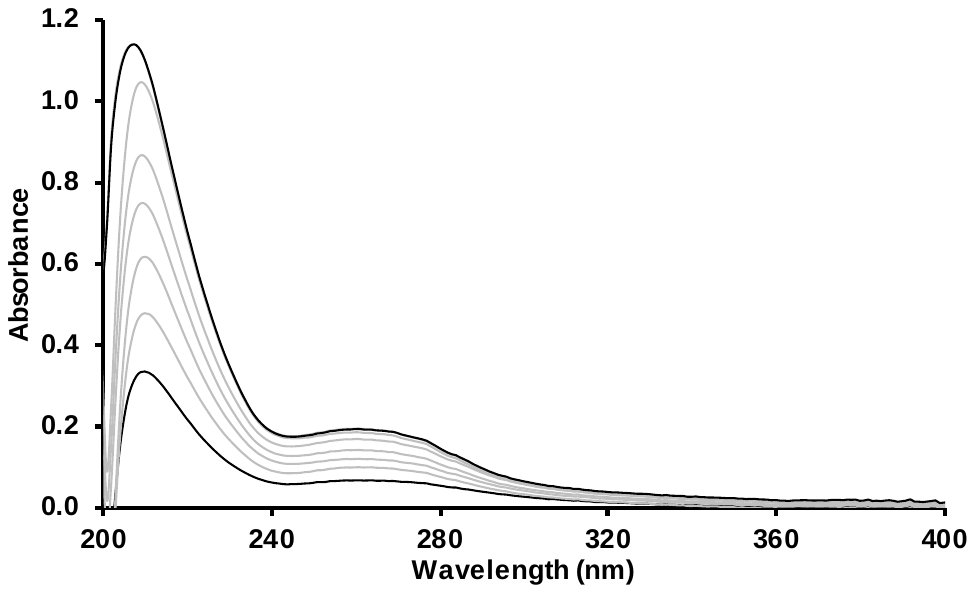

**b**

| Exposure time (hour) | Absorbance |
| --- | --- |
| 0 | 0 |
| 1 | 0.0639915 |
| 2 | 0.098074 |
| 3 | 0.118259 |
| 4 | 0.139254 |
| 5 | 0.16778 |
| 6 | 0.184645 |
| 7 | 0.190691 |
| 8 | 0.193314 |

Figure S9. Daylight-mediated lifitegrast release from a contact lens (Trial - 1). (a) a lifitegrast release contact lens was immersed in 2 mL of PBS placed on a stir plate at a rate of 60 rpm, and exposed to indoor daylight. 1 mL of the bathing solution was withdrawn, and then returned to the mother solution after its UV-vis spectra was recorded every hour, (b) plot of the absorption at 260 nm versus time.

**a**

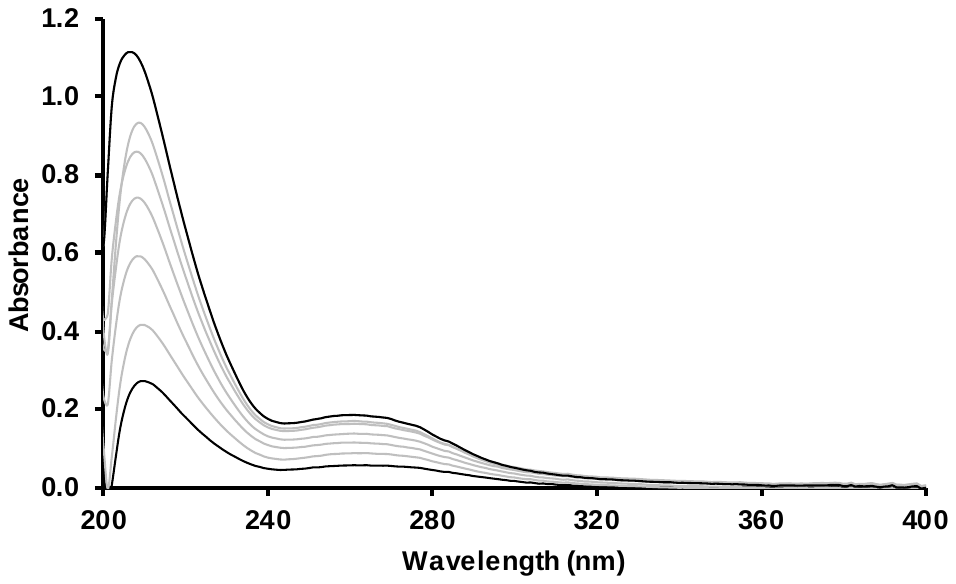

**b**

| Exposure time (hour) | Absorbance |
| --- | --- |
| 0 | 0 |
| 1 | 0.058115 |
| 2 | 0.086628 |
| 3 | 0.115309 |
| 4 | 0.136467 |
| 5 | 0.161324 |
| 6 | 0.170242 |
| 7 | 0.186354 |
| 8 | 0.186354 |

Figure S10. Daylight-mediated lifitegrast release from a contact lens (Trial - 2). (a) a lifitegrast release contact lens was immersed in 2 mL of PBS, placed on a stir plate at a rate of 60 rpm, and exposed to indoor daylight. 1 mL of the bathing solution was withdrawn, and then returned to the mother solution after its UV-vis spectra was recorded every hour, (b) plot of the absorption at 260 nm versus time.

**a**

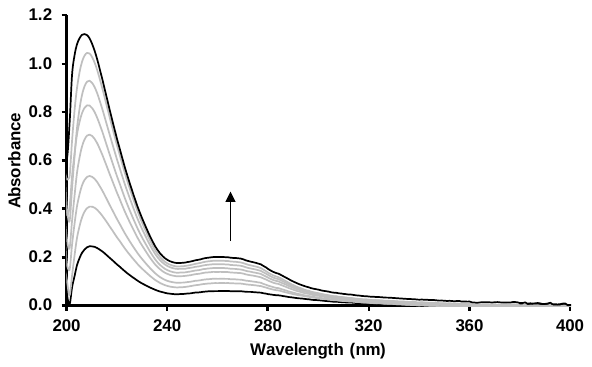

**b**

| Exposure time (hour) | Absorbance |
| --- | --- |
| 0 | 0 |
| 1 | 0.0607128 |
| 2 | 0.0928783 |
| 3 | 0.111301 |
| 4 | 0.139126 |
| 5 | 0.156067 |
| 6 | 0.172337 |
| 7 | 0.186922 |
| 8 | 0.201487 |

Figure S11. Daylight-mediated lifitegrast release from a contact lens (Trial - 3). (a) a lifitegrast release contact lens was immersed in 2 mL of PBS, placed on a stir plate at a rate of 60 rpm, and exposed to indoor daylight. 1 mL of the bathing solution was withdrawn, and then returned to the mother solution after its UV-vis spectra was recorded every hour, (b) plot of the absorption at 260 nm versus time.

| Exposure time (hour) | Concentration (µM) |
| --- | --- |
| 0 | 0 |
| 1 | 5.3929 |
| 2 | 8.18821 |
| 3 | 10.1731 |
| 4 | 12.2374 |
| 5 | 14.3118 |
| 6 | 15.5523 |
| 7 | 16.6362 |
| 8 | 17.1432 |

Figure S12. A graph of the indoor daylight mediated lifitegrast release (n = 3) from a contact lens, including the average and standard deviation.

### **(5) High-resolution Electrospray Ionization Mass Spectrometry**

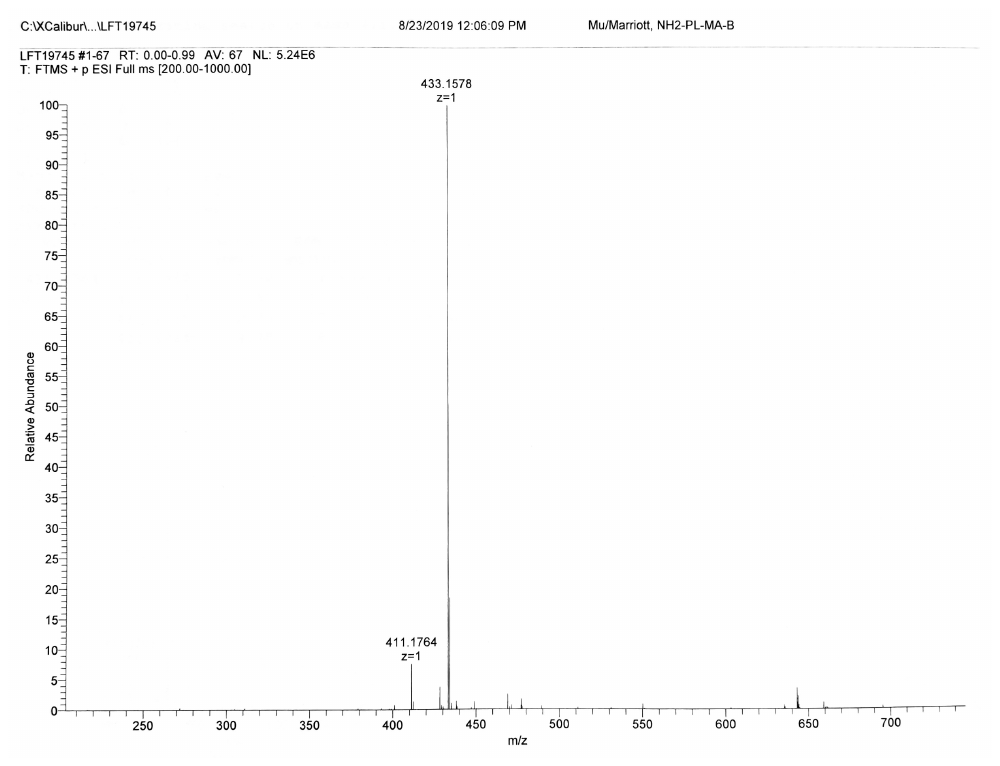

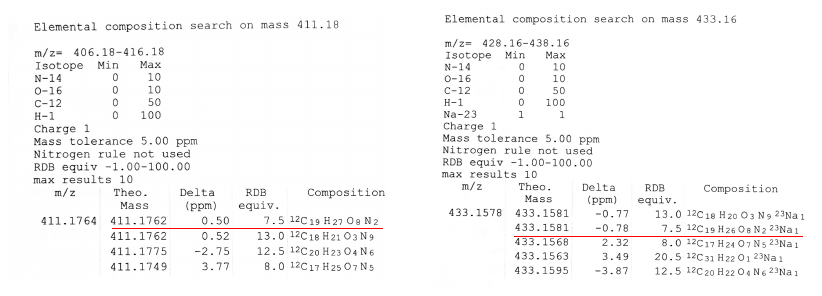

Figure S13. High-resolution electrospray ionization mass spectrometry (HR ESI-MS) of **PL-MA** m/z calculated for C_19_H_27_O_8_N_2_ ([M+H]^+^) 411.1762, found 411.1764, delta 0.50 ppm; m/z calculated for C_19_H_26_O_8_N_2_Na_1_ ([M+Na]^+^) 433.1581, found 433.1578, delta -0.78 ppm.

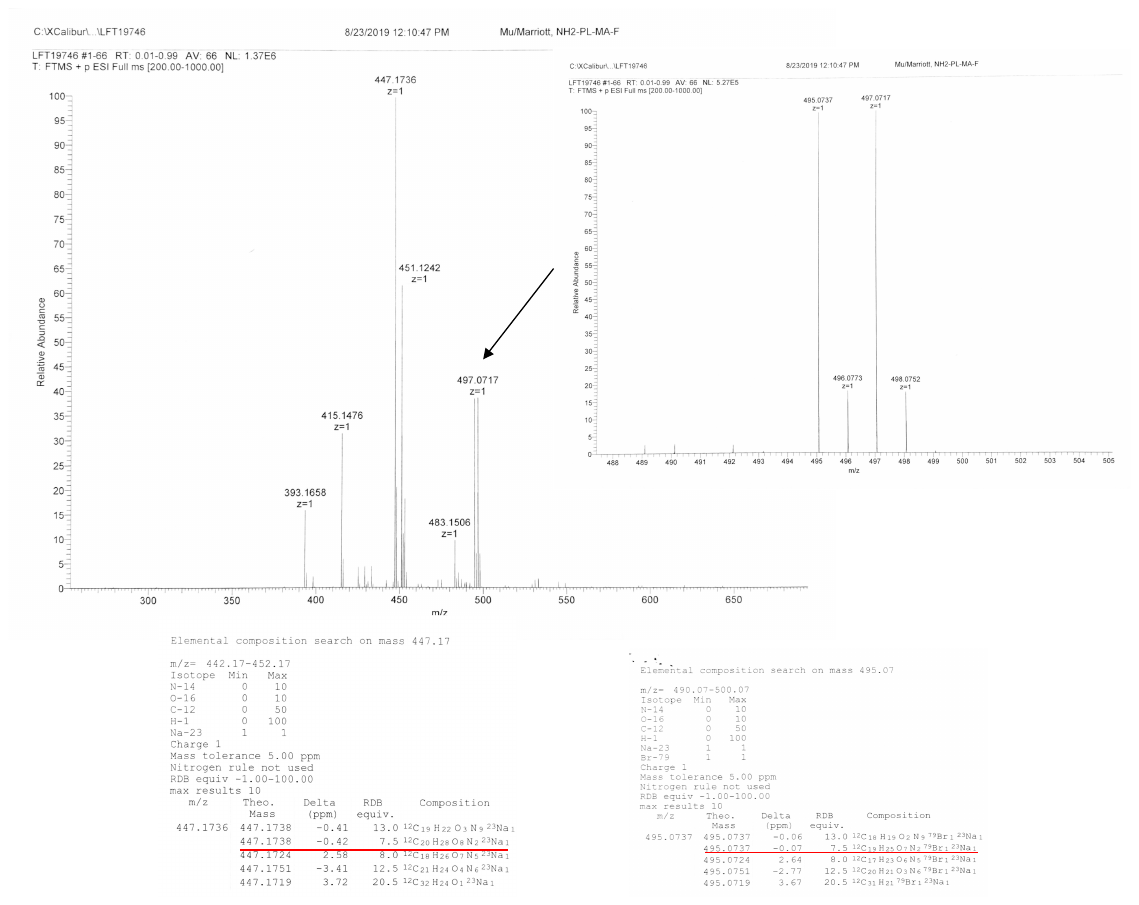

Figure S14. HR ESI-MS of **Br-PL-MA** m/z calculated for C_19_H_25_O_7_N_2_Br_1_Na_1_ ([M+Na]^+^) 495.0737, found 495.0737, delta -0.07 ppm (insert). The signal found at 447.1736 was an product from an unexpected reaction between **Br-PL-MA** with MeOH solvent during the procedure of MS measurement, calculated for C_20_H_28_O_8_N_2_Na_1_ ([M+Ma]^+^) 447.1744

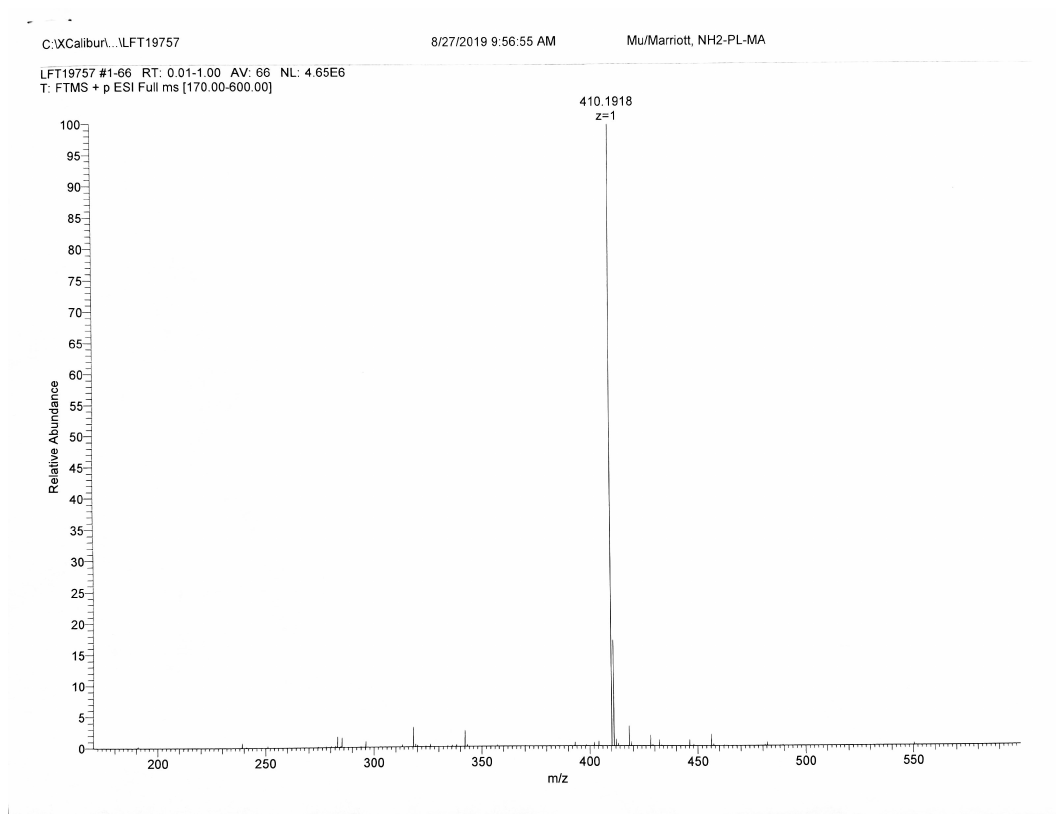

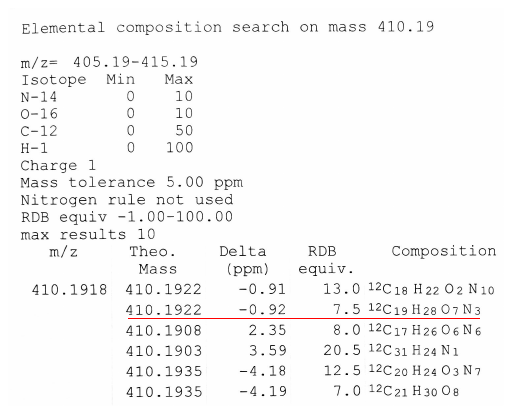

Figure S15. HR ESI-MS of **NH_2_-PL-MA** m/z calculated for C_19_H_28_O_7_N_3_ ([M+H]^+^) 410.1922, found 410.1918, delta -0.92 ppm.

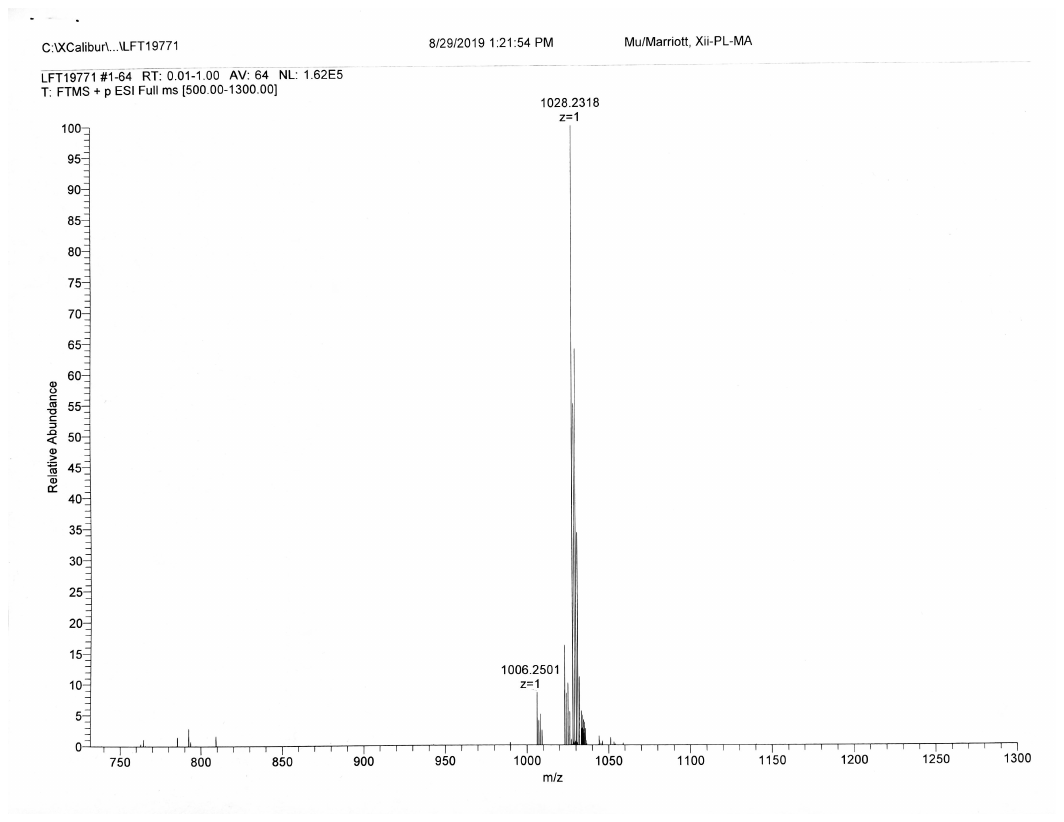

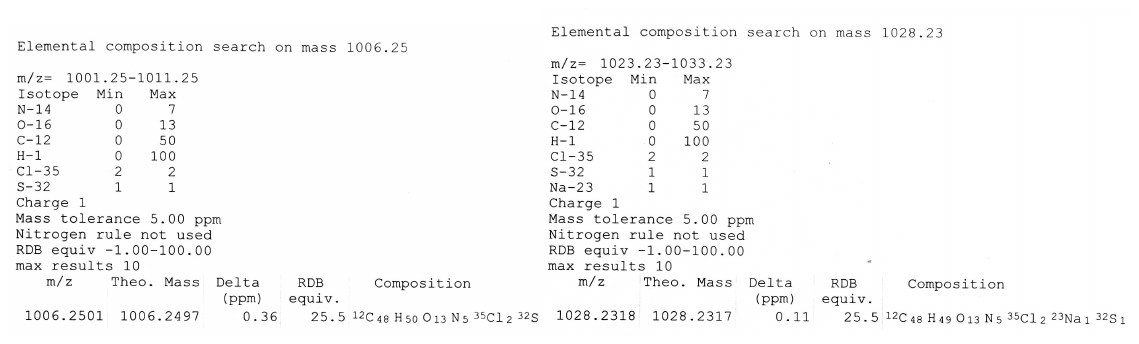

Figure S16. HR ESI-MS of **LG-PL-MA** m/z calculated for C_48_H_50_O_13_N_5_Cl_2_S_1_ ([M+H]^+^) 1006.2497, found 1006.2501, delta 0.36 ppm; m/z calculated for C_48_H_49_O_13_N_5_Cl_2_Na_1_S_1_ ([M+Na]^+^) 1028.2317, found 1028.2318, delta 0.11 ppm.

### **(6) UV-vis absorption spectra of synthetic compounds**

Absorbance

| Wavelength (nm) | Absorbance (AU) |
| --- | --- |
| 190 | 1.09619 |
| 191 | 0.389348 |
| 192 | 0.482857 |
| 193 | 0.559432 |
| 194 | 0.680745 |
| 195 | 0.481092 |
| 196 | 0.953854 |
| 197 | 0.963202 |
| 198 | 0.810455 |
| 199 | 1.06642 |
| 200 | 0.963959 |
| 201 | 0.576188 |
| 202 | 0.77905 |
| 203 | 0.680384 |
| 204 | 0.756258 |
| 205 | 0.782517 |
| 206 | 0.83256 |
| 207 | 0.698607 |
| 208 | 0.918422 |
| 209 | 1.01325 |
| 210 | 0.777597 |
| 211 | 0.74309 |
| 212 | 0.901968 |
| 213 | 0.943966 |
| 214 | 0.91617 |
| 215 | 0.924935 |
| 216 | 0.941403 |
| 217 | 0.947876 |
| 218 | 0.871241 |
| 219 | 0.929413 |
| 220 | 0.935063 |
| 221 | 0.89826 |
| 222 | 0.929587 |
| 223 | 0.956496 |
| 224 | 0.96429 |
| 225 | 0.953439 |
| 226 | 0.984637 |
| 227 | 1.03985 |
| 228 | 1.07601 |
| 229 | 1.09392 |
| 230 | 1.08364 |
| 231 | 1.05717 |
| 232 | 1.03252 |
| 233 | 1.01062 |
| 234 | 0.994526 |
| 235 | 0.985955 |
| 236 | 0.981385 |
| 237 | 0.981887 |
| 238 | 0.986849 |
| 239 | 0.994582 |
| 240 | 1.00617 |
| 241 | 1.01969 |
| 242 | 1.0357 |
| 243 | 1.05145 |
| 244 | 1.06634 |
| 245 | 1.07784 |
| 246 | 1.08322 |
| 247 | 1.08212 |
| 248 | 1.07384 |
| 249 | 1.0582 |
| 250 | 1.03689 |
| 251 | 1.01151 |
| 252 | 0.979551 |
| 253 | 0.94144 |
| 254 | 0.894179 |
| 255 | 0.837194 |
| 256 | 0.772201 |
| 257 | 0.702016 |
| 258 | 0.629417 |
| 259 | 0.559907 |
| 260 | 0.496704 |
| 261 | 0.443333 |
| 262 | 0.398861 |
| 263 | 0.363841 |
| 264 | 0.33753 |
| 265 | 0.318774 |
| 266 | 0.30537 |
| 267 | 0.296296 |
| 268 | 0.291047 |
| 269 | 0.288039 |
| 270 | 0.286971 |
| 271 | 0.287396 |
| 272 | 0.28879 |
| 273 | 0.291012 |
| 274 | 0.294282 |
| 275 | 0.298691 |
| 276 | 0.303572 |
| 277 | 0.309979 |
| 278 | 0.31706 |
| 279 | 0.32459 |
| 280 | 0.332855 |
| 281 | 0.341554 |
| 282 | 0.350344 |
| 283 | 0.359265 |
| 284 | 0.36783 |
| 285 | 0.376424 |
| 286 | 0.384556 |
| 287 | 0.392552 |
| 288 | 0.400293 |
| 289 | 0.407613 |
| 290 | 0.414306 |
| 291 | 0.420736 |
| 292 | 0.426875 |
| 293 | 0.432391 |
| 294 | 0.437502 |
| 295 | 0.442137 |
| 296 | 0.446148 |
| 297 | 0.449867 |
| 298 | 0.453342 |
| 299 | 0.456013 |
| 300 | 0.458132 |
| 301 | 0.460202 |
| 302 | 0.461714 |
| 303 | 0.462945 |
| 304 | 0.463565 |
| 305 | 0.464119 |
| 306 | 0.464113 |
| 307 | 0.464167 |
| 308 | 0.463469 |
| 309 | 0.462628 |
| 310 | 0.462424 |
| 311 | 0.461478 |
| 312 | 0.460771 |
| 313 | 0.459662 |
| 314 | 0.458571 |
| 315 | 0.457634 |
| 316 | 0.456318 |
| 317 | 0.455191 |
| 318 | 0.454387 |
| 319 | 0.453453 |
| 320 | 0.452838 |
| 321 | 0.452735 |
| 322 | 0.45292 |
| 323 | 0.453542 |
| 324 | 0.454654 |
| 325 | 0.456365 |
| 326 | 0.458215 |
| 327 | 0.460517 |
| 328 | 0.463216 |
| 329 | 0.46632 |
| 330 | 0.469786 |
| 331 | 0.472902 |
| 332 | 0.476256 |
| 333 | 0.479578 |
| 334 | 0.482638 |
| 335 | 0.485751 |
| 336 | 0.488514 |
| 337 | 0.491068 |
| 338 | 0.493483 |
| 339 | 0.495886 |
| 340 | 0.498472 |
| 341 | 0.500529 |
| 342 | 0.502242 |
| 343 | 0.503711 |
| 344 | 0.505189 |
| 345 | 0.506335 |
| 346 | 0.506789 |
| 347 | 0.507323 |
| 348 | 0.507204 |
| 349 | 0.506345 |
| 350 | 0.505647 |
| 351 | 0.503855 |
| 352 | 0.501555 |
| 353 | 0.499475 |
| 354 | 0.496291 |
| 355 | 0.492877 |
| 356 | 0.489092 |
| 357 | 0.485377 |
| 358 | 0.481003 |
| 359 | 0.47666 |
| 360 | 0.472315 |
| 361 | 0.466473 |
| 362 | 0.460878 |
| 363 | 0.454285 |
| 364 | 0.447458 |
| 365 | 0.440118 |
| 366 | 0.43272 |
| 367 | 0.424558 |
| 368 | 0.415739 |
| 369 | 0.406942 |
| 370 | 0.39749 |
| 371 | 0.388286 |
| 372 | 0.378677 |
| 373 | 0.36906 |
| 374 | 0.359714 |
| 375 | 0.349797 |
| 376 | 0.339752 |
| 377 | 0.329656 |
| 378 | 0.319551 |
| 379 | 0.309706 |
| 380 | 0.299564 |
| 381 | 0.290274 |
| 382 | 0.279872 |
| 383 | 0.269143 |
| 384 | 0.258302 |
| 385 | 0.247749 |
| 386 | 0.236944 |
| 387 | 0.226141 |
| 388 | 0.215662 |
| 389 | 0.205491 |

Figure S17. UV-vis absorption spectrum of **NH_2_-PL-MA** in MeOH with three absorption peaks at 250, 300, and 350 nm.

| Wavelength (nm) | Absorbance |
| --- | --- |
| 190 | 0.733946 |
| 191 | 0.784157 |
| 192 | 1.1325 |
| 193 | 1.01255 |
| 194 | 0.769516 |
| 195 | 0.896906 |
| 196 | 0.784518 |
| 197 | 1.06712 |
| 198 | 1.26656 |
| 199 | 1.08803 |
| 200 | 1.41414 |
| 201 | 1.51978 |
| 202 | 1.81611 |
| 203 | 1.76706 |
| 204 | 2.39719 |
| 205 | 2.56663 |
| 206 | 2.41097 |
| 207 | 2.54231 |
| 208 | 3.02541 |
| 209 | 2.89699 |
| 210 | 2.78623 |
| 211 | 2.95442 |
| 212 | 2.98953 |
| 213 | 3.0174 |
| 214 | 3.01637 |
| 215 | 2.96134 |
| 216 | 2.85239 |
| 217 | 2.85892 |
| 218 | 2.81026 |
| 219 | 2.77335 |
| 220 | 2.68394 |
| 221 | 2.62563 |
| 222 | 2.54893 |
| 223 | 2.44659 |
| 224 | 2.33864 |
| 225 | 2.24997 |
| 226 | 2.14608 |
| 227 | 2.03244 |
| 228 | 1.92536 |
| 229 | 1.82561 |
| 230 | 1.72481 |
| 231 | 1.63509 |
| 232 | 1.55078 |
| 233 | 1.46274 |
| 234 | 1.3793 |
| 235 | 1.29945 |
| 236 | 1.22658 |
| 237 | 1.16265 |
| 238 | 1.11104 |
| 239 | 1.07124 |
| 240 | 1.04099 |
| 241 | 1.01978 |
| 242 | 1.00592 |
| 243 | 0.997102 |
| 244 | 0.99299 |
| 245 | 0.991573 |
| 246 | 0.991619 |
| 247 | 0.993879 |
| 248 | 0.996612 |
| 249 | 0.998693 |
| 250 | 1.00037 |
| 251 | 1.00131 |
| 252 | 1.0012 |
| 253 | 0.999785 |
| 254 | 0.996149 |
| 255 | 0.990771 |
| 256 | 0.980569 |
| 257 | 0.967727 |
| 258 | 0.952943 |
| 259 | 0.938179 |
| 260 | 0.92317 |
| 261 | 0.910377 |
| 262 | 0.898508 |
| 263 | 0.885265 |
| 264 | 0.871676 |
| 265 | 0.858351 |
| 266 | 0.84583 |
| 267 | 0.835454 |
| 268 | 0.825715 |
| 269 | 0.812401 |
| 270 | 0.795117 |
| 271 | 0.773259 |
| 272 | 0.750539 |
| 273 | 0.730125 |
| 274 | 0.715103 |
| 275 | 0.704025 |
| 276 | 0.692497 |
| 277 | 0.674454 |
| 278 | 0.650145 |
| 279 | 0.620686 |
| 280 | 0.587068 |
| 281 | 0.554311 |
| 282 | 0.525259 |
| 283 | 0.504434 |
| 284 | 0.489851 |
| 285 | 0.47313 |
| 286 | 0.450259 |
| 287 | 0.422582 |
| 288 | 0.393823 |
| 289 | 0.365455 |
| 290 | 0.338374 |
| 291 | 0.312593 |
| 292 | 0.288197 |
| 293 | 0.266763 |
| 294 | 0.247766 |
| 295 | 0.231138 |
| 296 | 0.216739 |
| 297 | 0.204723 |
| 298 | 0.194471 |
| 299 | 0.185595 |
| 300 | 0.177913 |
| 301 | 0.171362 |
| 302 | 0.165486 |
| 303 | 0.160134 |
| 304 | 0.155196 |
| 305 | 0.150668 |
| 306 | 0.146423 |
| 307 | 0.14186 |
| 308 | 0.137838 |
| 309 | 0.134059 |
| 310 | 0.131047 |
| 311 | 0.127999 |
| 312 | 0.125149 |
| 313 | 0.122139 |
| 314 | 0.119857 |
| 315 | 0.117426 |
| 316 | 0.115016 |
| 317 | 0.113012 |
| 318 | 0.111223 |
| 319 | 0.109392 |
| 320 | 0.107859 |
| 321 | 0.106755 |
| 322 | 0.105688 |
| 323 | 0.104685 |
| 324 | 0.10402 |
| 325 | 0.103451 |
| 326 | 0.103094 |
| 327 | 0.102695 |
| 328 | 0.102366 |
| 329 | 0.102097 |
| 330 | 0.102044 |
| 331 | 0.102059 |
| 332 | 0.101933 |
| 333 | 0.101787 |
| 334 | 0.101717 |
| 335 | 0.101586 |
| 336 | 0.101659 |
| 337 | 0.101461 |
| 338 | 0.101259 |
| 339 | 0.10113 |
| 340 | 0.100918 |
| 341 | 0.100904 |
| 342 | 0.100679 |
| 343 | 0.100478 |
| 344 | 0.100051 |
| 345 | 0.0994391 |
| 346 | 0.0990715 |
| 347 | 0.0985713 |
| 348 | 0.097815 |
| 349 | 0.0969324 |
| 350 | 0.0959959 |
| 351 | 0.0953283 |
| 352 | 0.0945463 |
| 353 | 0.0931787 |
| 354 | 0.0922704 |
| 355 | 0.0907984 |
| 356 | 0.0895362 |
| 357 | 0.0883684 |
| 358 | 0.0873609 |
| 359 | 0.0863299 |
| 360 | 0.0850649 |
| 361 | 0.0835152 |
| 362 | 0.0822029 |
| 363 | 0.0806365 |
| 364 | 0.0787497 |
| 365 | 0.076807 |
| 366 | 0.0751114 |
| 367 | 0.07412 |
| 368 | 0.0723472 |
| 369 | 0.0704513 |
| 370 | 0.068347 |
| 371 | 0.0663285 |
| 372 | 0.0646992 |
| 373 | 0.062706 |
| 374 | 0.0600224 |
| 375 | 0.059278 |
| 376 | 0.057776 |
| 377 | 0.0552173 |
| 378 | 0.0542412 |
| 379 | 0.0543489 |
| 380 | 0.0520334 |
| 381 | 0.0494165 |
| 382 | 0.0471721 |
| 383 | 0.0463371 |
| 384 | 0.0441585 |
| 385 | 0.0423484 |
| 386 | 0.0397959 |
| 387 | 0.0390558 |
| 388 | 0.0405092 |
| 389 | 0.0385447 |

Figure S18. UV-vis absorption spectrum of **LG-PL-MA** in MeOH with two absorption peaks at 215 and 260 nm.
